# Cell phenotypes in the biomedical literature: a systematic analysis and text mining corpus

**DOI:** 10.64898/2026.02.11.705457

**Authors:** Noam H. Rotenberg, Robert Leaman, Rezarta Islamaj, Helena Kuivaniemi, Gerard Tromp, Brian Fluharty, Savannah Richardson, Caroline Eastwood, Matthew Diller, Bingfang Xu, Ajith V. Pankajam, David Osumi-Sutherland, Zhiyong Lu, Richard H. Scheuermann

**Author notes:** These authors contributed equally to this work (Noam H. Rotenberg, Robert Leaman).

## Abstract

The variety of cell phenotypes identified by single-cell technologies is rapidly expanding, yet this knowledge is dispersed across the scientific literature and incompletely represented in structured resources. We present the CellLink corpus, a manually annotated collection of over 22,000 mentions of human and mouse cell populations in recent journal articles, distinguishing specific cell phenotypes, heterogeneous cell populations, and vague cell populations, and linking to Cell Ontology (CL) terms as either exact or related matches, covering nearly half of the terms in the current CL. A systematic analysis reveals lineage-specific patterns in how authors utilize anatomical context, molecular signatures, functional roles, developmental stage, and other attributes in cell naming. We show that fine-tuning transformer-based models on CellLink yields strong performance for named entity recognition, while embedding-based approaches support zero-shot entity linking and distinguishing exact from related matches. We further demonstrate the utility of CellLink to expand and refine the chondrocyte branch of CL.

## Introduction

Advances in single-cell omics technologies are transforming our understanding of cell biology^1, 2^. These methods enable analyses of cell phenotypes (a cell’s expressed type and state) by evaluating gene expression, chromatin structure, and spatial context at single-cell resolution. As a result, researchers are frequently identifying, defining, and naming novel cell phenotypes based on single-cell experimental data^1, 2^. For example, single-cell RNA sequencing (scRNA-seq) experiments can identify novel cell phenotypes by the specific combinations of gene transcripts they express, their so-called transcriptional profile. The Human Cell Atlas^2^, HuBMAP^3^, CellxGene^4^, the Cell Ontology (CL)^5^ and other projects capture and curate many of these emerging cell phenotypes.

The rapid adoption of single-cell technologies has not only accelerated discovery but also shifted how researchers describe cell phenotypes in the literature. Publications mentioning scRNA-seq and related cell typing methods have risen dramatically in recent years (Fig. 1a, 1b), and this increase has been accompanied by a marked shift in terminology (Fig. 1c). Earlier work more frequently emphasized molecular and signaling mechanisms studied in cell lines or experimental model systems. In contrast, more recent studies focus on scRNA-seq-based transcriptomic analyses of patient-derived samples, highlighting heterogeneity in gene expression signatures, immune and tumor microenvironments, and associations with disease progression and clinical outcomes. This shift reflects the increasing use of computational methods to analyze large single-cell datasets across cohorts, enabling integrative and clinically relevant characterization of cellular populations.

**Fig. 1.**
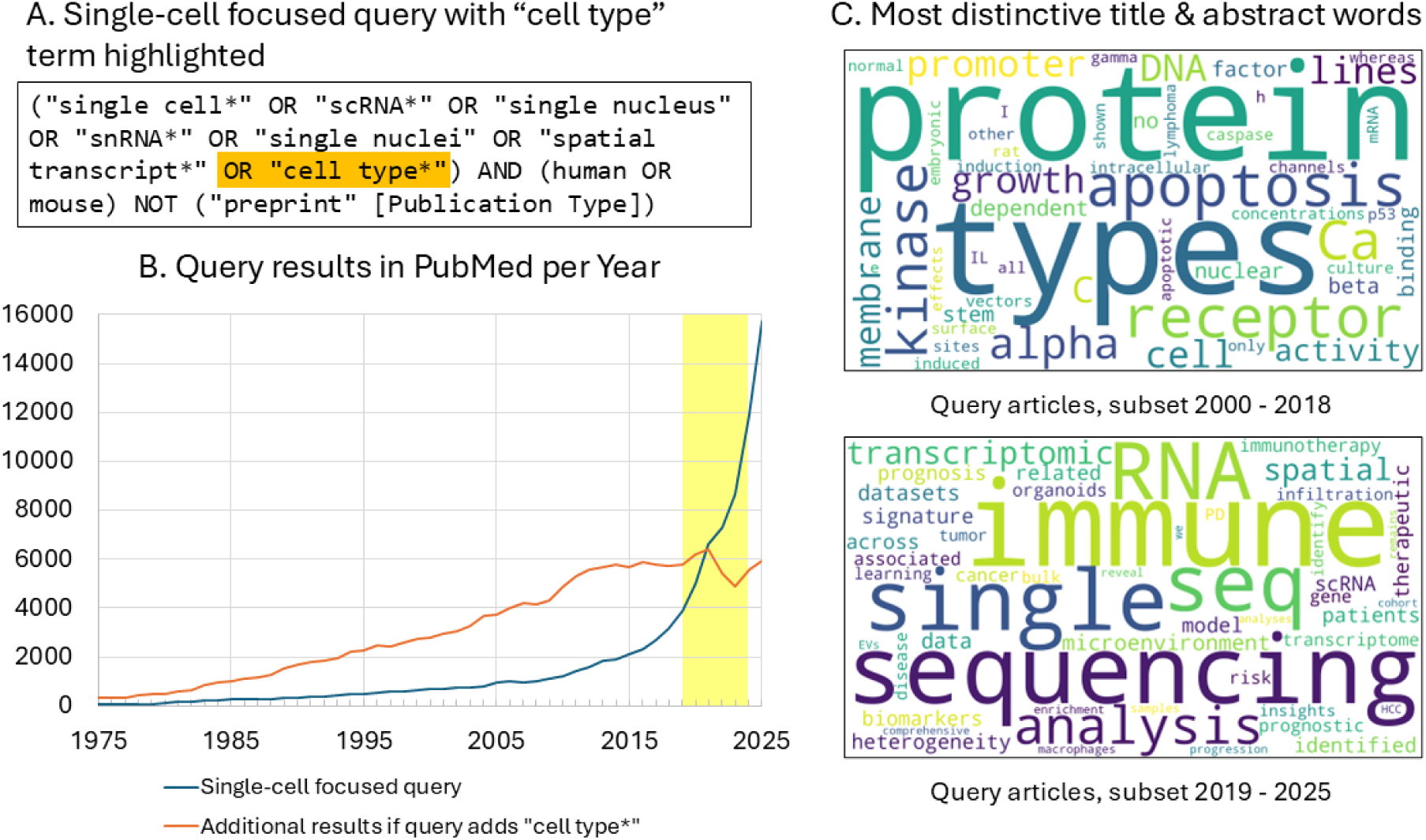
Growth of single cell genomics and its impact on cell type terminology in the biomedical literature. (A) PubMed query focused on single-cell studies in human and mouse, with an expansion (orange) including the term “cell type.” (B) Annual number of PubMed results retrieved by the single-cell-focused query alone (blue) and the additional results retrieved when “cell type” is included (orange). The region shaded yellow marks the time period focused on in this study (2019–2024). The query was executed on Jan 14, 2026. (C) Words enriched in titles and abstracts of retrieved articles across two time periods, (2000–2018 vs. 2019–2025). Enrichment was quantified using informative log-odds ratios with a Dirichlet prior^6^, contrasting each period with the other. Positive z-scores were used as weights to generate word clouds with the wordcloud Python library (https://github.com/amueller/word_cloud).

Cell phenotypes are described in diverse ways, using marker genes, tissue of origin, developmental trajectory, functional role, and cell state. These descriptive strategies reflect a rapidly expanding field in which overlapping phenotypes may be labeled inconsistently, and new terms may not yet be connected to established concepts. For biologists seeking to locate and compare relevant studies, this variability can obscure important findings and complicate interpretation.

Computational methods such as natural language processing (NLP) have been widely used to help researchers search, organize, and synthesize the biomedical literature^7–13^. NLP systems often use machine learning models trained on annotated text corpora; these corpora provide examples of how biological entities of interest are expressed in the text. Some off-the-shelf models do not require such datasets for training, but they do require annotated text data for system development and evaluation. However, when terminology evolves rapidly – as with the rise of scRNA-seq – NLP systems prepared using older corpora often fail to recognize new or unconventional expressions, limiting their effectiveness for search, synthesis, and curation. As a result, important, novel results may remain hidden. Robust training and evaluation of these systems requires resources that capture the current language of cell phenotypes in the literature.

Several existing named entity corpora (annotated text datasets) include cell population annotations, including the Anatomical Entity Mention (AnatEM) corpus^14^, the BioCreative VI BioID corpus^15^, the Colorado Richly Annotated Full Text (CRAFT) corpus^16^, and the corpus used for the bioentity recognition task at the 2004 Joint Workshop on Natural Language Processing in Biomedicine and its Applications (JNLPBA)^17^. However, all were created before the widespread adoption of robust single-cell transcriptomic methods and therefore do not reflect the variety and granularity of cell phenotypes being described in more recent literature, particularly those derived from scRNA-seq studies in human tissues and mouse models. Moreover, these datasets were not designed to focus on cell phenotypes: CRAFT focuses on mouse genetics, while JNLPBA is derived from the GENIA corpus^18^, which focuses on blood transcription factors. Consequently, their coverage of cell phenotypes is limited in both breadth and depth, with relatively few instances of the fine-grained phenotypes discussed in the recent scRNA-seq literature. Finally, only BioID and CRAFT link their cell phenotype annotations to identifiers from structured resources such as CL, and the combined total number of distinct identifiers is small (n=378). These limitations reduce their utility for developing and evaluating NLP systems to identify the expanding set of cell phenotypes described in the current literature.

In this work, we introduce the National Library of Medicine (NLM) CellLink corpus, a new set of annotated excerpts from full-text biomedical articles published between 2019 and 2024. The corpus was manually annotated by experienced biologists for mentions of human and mouse cell phenotypes, focusing on naturally occurring cells rather than cells that have been experimentally modified. The goals of this work are to (1) catalogue and characterize how authors refer to cell phenotypes in recent biomedical publications; (2) provide training and evaluation data for informatics systems that can recognize both established and novel cell types; and (3) identify novel cell types reported in the recent literature and demonstrate their integration into the Cell Ontology through a curator-led case study. This work also establishes a foundation for future efforts to create a knowledgebase of extracted relationships between cell types and other key biomedical entities, such as marker genes, anatomical structures, perturbation responses, and disease states.

## Results

### Construction and overview of the CellLink corpus

The CellLink corpus was developed to systematically sample how authors describe cell phenotypes in the recent biomedical literature. Articles were selected from PubMed (2019−2024) using a query emphasizing single-cell and related studies in human and mouse, ensuring that the articles returned were both biomedically relevant and reflected the recent vocabulary shift toward high resolution cell characterization (Fig. 1). Passages – either paragraphs or equivalent textual units from full text scientific journal articles if available, otherwise the title and abstract – were chosen as the annotation unit to balance context with annotation efficiency.

Passages were selected for annotation using a sampling strategy designed to balance representativeness with enrichment of less frequent characteristics. Specifically, we represent each passage from the query articles by a rich set of discrete characteristics and their corresponding values, including publication year, passage length, and the count of each word present. We also defined a set of clusters of Medical Subject Headings (MeSH) terms representing anatomical systems (e.g., “Nervous-Anatomy”) or diseases of a specific system (e.g., “Cardiovascular-Disease”); the full list is shown in Supplementary Table 1. Passages were then selected incrementally by matching the distribution of characteristics in the selected set to that of all query passages (the target distribution). Prior to selection, the target distribution was adjusted to (1) increase the sampling of infrequent characteristics, (2) increase the sampling of characteristics related to our focus, and (3) decrease the sampling of characteristics outside our focus, such as cell lines and cancers. The passage selection method is described in detail in Methods: Article retrieval and passage selection.

The distribution of the MeSH clusters before and after designed sampling is shown in Supplementary Table 2. Passages from articles discussing the anatomy of the immune and nervous systems were the most overrepresented in the corpus, which is consistent with the annotators reporting that passages discussing these cell types had high complexity. Passages from articles discussing the exocrine system were the most relatively overrepresented, as they were the rarest in the original distribution (0.7% prevalence). Passages with cell lines and cancer MeSH categories were selectively under-sampled, as designed, to focus on healthy, naturally occurring cells.

Four experienced cell biologists read and annotated all passages following detailed guidelines (see Supplementary File 1). Preliminary analysis revealed that many of the cell populations mentioned in the literature do not refer to discrete, identifiable cell phenotypes. The annotators therefore classified the text spans they identified as referring to cell populations as one of three types: cell phenotype, heterogeneous cell population, or vague cell population. A cell phenotype is distinct, identifiable cell type or state, for example, “endothelial cells”, “oligodendrocytes”, or “exhausted CD8+ T cells”. A heterogeneous cell population refers to a group of cells from unrelated developmental lineages, typically characterized by a shared attribute, for example: “kidney cells”, “somatic cells”, or “tumor cells”. A vague cell population refers to a group of cells whose identities cannot be determined from the text provided; linguistically, these are often descriptions rather than names, for example: “T-cell populations”, “myeloid cell clusters”, or “non-neuronal cells”, “HO1-like cells”. Note that parts of these phrases, such as “T-cell”, could additionally be annotated as cell phenotypes or heterogeneous cell populations. These entity types are described in detail in the annotation guidelines (Supplementary File 1). Cell phenotype and heterogeneous cell population mentions were linked to identifiers in the Cell Ontology according to their meaning in context and labeled as either an exact or related match (Fig. 2). Annotators marked matches as exact when the term meaning was equivalent, or related when the linked term was broader, narrower, or otherwise contextually associated. Additional examples are available in Supplementary Table 3.

**Fig. 2.**
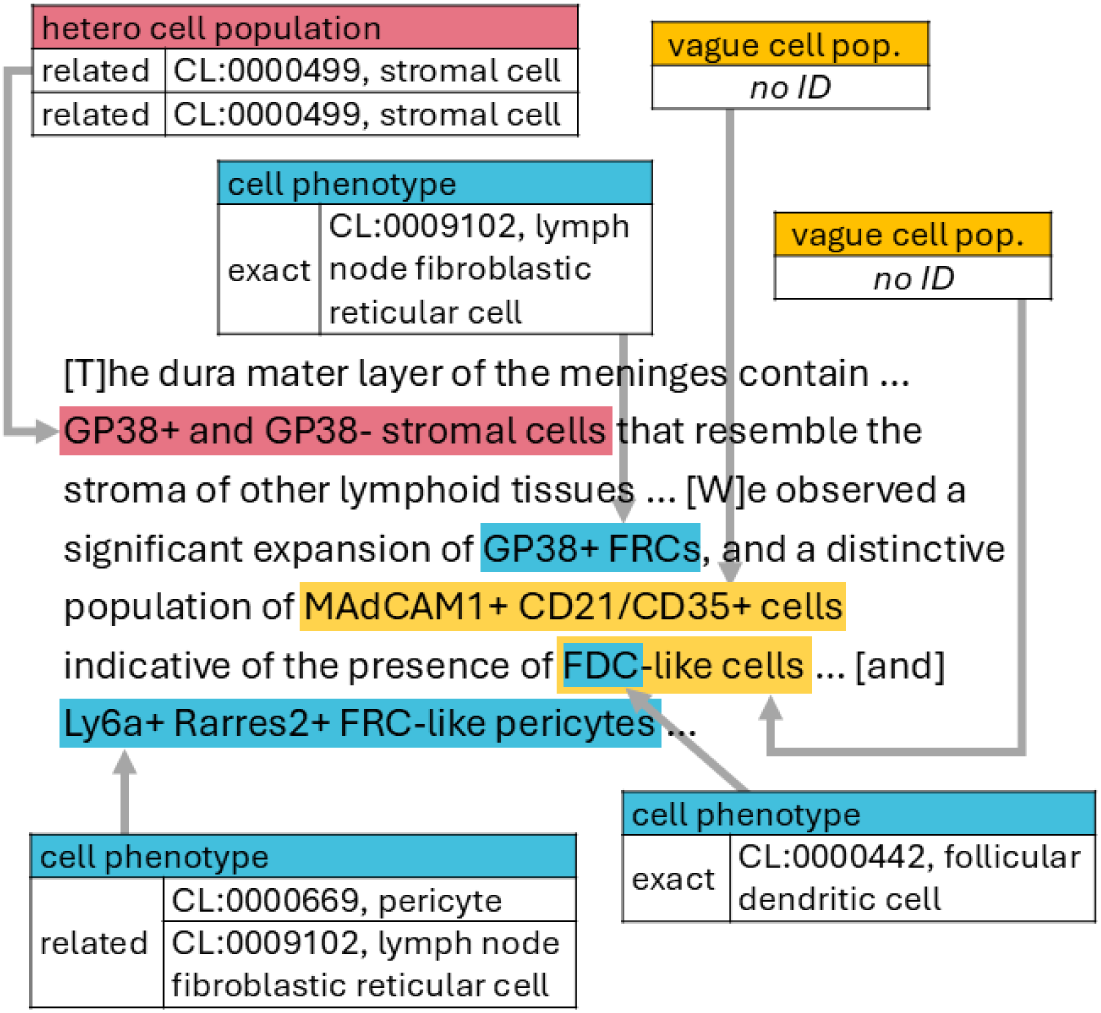
Example of CellLink annotations. Excerpt from the CellLink corpus illustrating annotations for cell phenotypes, heterogeneous cell populations, and vague cell populations. Cell phenotype and heterogeneous cell population annotations are linked to Cell Ontology (CL) identifiers and labeled as exact or related matches; vague cell populations are not linked. Example simplified from PMID 37983289. FDC, follicular dendritic cell; FRC, fibroblastic reticular cell.

Each passage was independently annotated by two annotators following a three-round process described previously^19^. In the first round, annotators worked fully independently; in the second round, they again worked independently but could view their partner’s annotations to assess discrepancies and inconsistencies. In the final round, remaining disagreements were discussed for collaborative resolution until consensus was reached. Annotation was performed over eight weeks, with the first two weeks reserved for training and revising the annotation guidelines. Following the training phase, the average inter-annotator agreement (IAA) in Round One was 0.69 (Supplementary Table 4), calculated based on exact matches of both the text span and any identifiers labeled as exact (identifiers labeled as related were excluded). IAA was highest for cell phenotypes, lower for heterogeneous cell populations, and lowest for vague cell populations. However, the overlapping-span IAA across all three entity types was 0.96 (merged labels), indicating that the annotators almost always agreed on the presence and general location of a cell population mention, but not necessarily the full extent of the phrase that captures its description. While IAA for most passage types was similar to the overall average, slightly lower agreement was observed in titles (0.60), section headers (0.62), and abstracts (0.66) (Supplementary Table 5).

The cumulative number of unique mentions and unique identifiers rose steadily throughout the annotation phase (Supplementary Fig. 1). The number of unique mentions rose roughly linearly for each of the mention types, with no clear indication of diminishing returns. In contrast, the number of unique identifiers for cell phenotype mentions linked exactly to CL followed a more concave trend, consistent with diminishing marginal discovery of CL identifiers as annotation progressed. Cell phenotypes linked as related matches and heterogeneous cell populations increased more slowly and variably. To ensure consistency of the final annotations between different passages, several rounds of inconsistency reviews were carried out before the consensus annotations of the entire corpus were finalized. The resulting corpus, CellLink, is a curated, high-coverage resource for both cell phenotype recognition and ontology linking.

CellLink contains 3,005 excerpts from 2,765 articles in 467 journals, totaling 338,445 tokens, including 20,541 unique tokens (case insensitive). Here, we defined tokens as contiguous sequences of letters or numbers and are roughly equivalent to words for the purpose of estimating corpus size. The corpus contains a total of 22,362 annotations. Of all annotations, 82.8% were labeled as cell phenotype, 7.2% as heterogeneous cell population, and 10.0% as vague cell population (Table 1). Annotation length varied substantially across types, with vague cell population mentions tending to be significantly longer than annotations of the other two types. Approximately 65% of the unique cell phenotype and 31% of the unique heterogeneous cell population mentions were linked to a CL identifier as an exact match. The remainder represent new or unreported cell types, most of which were linked to one or more CL identifiers as related matches. The distribution of annotation types varied by passage type: vague and heterogeneous cell populations were more frequent in titles, section headings, and abstracts, compared to cell phenotypes (Supplementary Table 5). In contrast, vague cell populations were particularly rare in tables, accounting for only 3% of annotations in those sections.

**Table 1.** Distribution and uniqueness of cell population annotations in CellLink*.

| Statistic | Cell phenotype | Heterogeneous cell population | Vague cell population | TOTAL<sup>†</sup> |
| --- | --- | --- | --- | --- |
| Number of cell population mentions | 18,515 (82.8%) | 1,616 (7.2%) | 2,231 (10.0%) | 22,362 (100.0%) |
| Unique mentions | 7,438 | 629 | 1,801 | 9,804 |
| Unique linked CL IDs | 1,214 | 231 | N/A | 1,251 |
| Unique mentions without exact CL ID | 2,573 | 435 | 1,801 | 4,783 |
| Annotation length<sup>‡</sup> (mean ± sd) | 13.5 ± 10.0 | 14.5 ± 8.6 | 23.9 ± 13.9 | 14.6 ± 10.8 |

\*The CellLink corpus comprises 3,005 excerpts containing 338,445 tokens (see Table 2).
<sup>†</sup>Note, a few unique mentions and identifiers overlap between entity types.
<sup>‡</sup>Number of characters
CL, Cell Ontology

The corpus was split into training (50.0%), validation (16.7%), and testing (33.3%) sets to support the training and testing of NLP methods for recognizing and linking cell phenotype mentions. The split was chosen to balance the number of annotations of each entity type, passage length, and the MeSH topic cluster frequency, while increasing the number of identifiers that only appear in a single set. The resulting subsets include many mentions not present in the other subsets. Supplementary Table 6 provides the distribution of various important features across the three subsets, while Supplementary Fig. 2 describes the number of unique mentions and identifiers shared across the three sets as Venn diagrams.

### Corpus validation and comparison

CellLink was compared with related corpora in terms of coverage and Named Entity Recognition (NER) model performance. We compared the CellLink corpus with four manually annotated corpora containing cell type annotations: AnatEM^14^, BioID^15^, CRAFT^16^, and JNLPBA^17^ (Table 2, additional details in Supplementary Table 7). Although most of these corpora contain more text overall, their broader annotation scope means that much of this text does not mention cell types. CellLink, however, is highly focused: it contains approximately the same number of total cell type annotations as the other corpora combined (22,362 vs. 22,292), but with nearly twice as many unique mentions (9,804 vs. 5,128) from only 13% of the number of sentences.

**Table 2.**
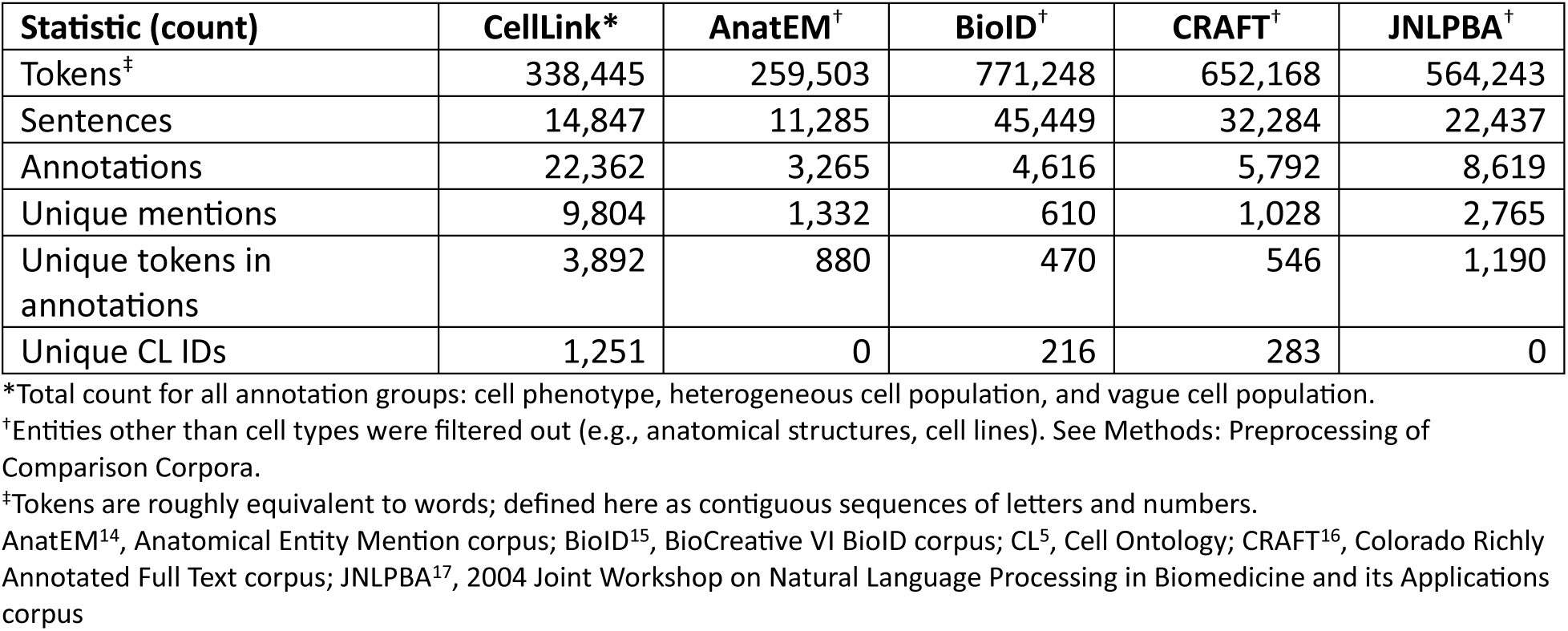
Comparison of CellLink with previous corpora containing cell type annotations.

| Statistic (count) | CellLink* | AnatEM<sup>†</sup> | BioID<sup>†</sup> | CRAFT<sup>†</sup> | JNLPBA<sup>†</sup> |
| --- | --- | --- | --- | --- | --- |
| Tokens<sup>‡</sup> | 338,445 | 259,503 | 771,248 | 652,168 | 564,243 |
| Sentences | 14,847 | 11,285 | 45,449 | 32,284 | 22,437 |
| Annotations | 22,362 | 3,265 | 4,616 | 5,792 | 8,619 |
| Unique mentions | 9,804 | 1,332 | 610 | 1,028 | 2,765 |
| Unique tokens in annotations | 3,892 | 880 | 470 | 546 | 1,190 |
| Unique CL IDs | 1,251 | 0 | 216 | 283 | 0 |

\*Total count for all annotation groups: cell phenotype, heterogeneous cell population, and vague cell population.
<sup>†</sup>Entities other than cell types were filtered out (e.g., anatomical structures, cell lines). See Methods: Preprocessing of Comparison Corpora.
<sup>‡</sup>Tokens are roughly equivalent to words; defined here as contiguous sequences of letters and numbers.
AnatEM<sup>14</sup>, Anatomical Entity Mention corpus; BioID<sup>15</sup>, BioCreative VI BioID corpus; CL<sup>5</sup>, Cell Ontology; CRAFT<sup>16</sup>, Colorado Richly Annotated Full Text corpus; JNLPBA<sup>17</sup>, 2004 Joint Workshop on Natural Language Processing in Biomedicine and its Applications corpus

Of the comparison datasets, only BioID and CRAFT contain links to CL, together covering only 378 unique CL identifiers. In contrast, CellLink contains a total of 1,251 unique identifiers, each labeled as either an exact or related match (Table 2). About one third of the unique mentions annotated as cell phenotypes (35%) were linked to a CL identifier with a related match or have no linkage, indicating that the cell type is either novel or currently unreported in CL.

CL is structured as a hierarchy with multiple levels of parent-child relations (a directed acyclic graph), where deeper nodes correspond to more specific cell types. CellLink provides much higher coverage of all major cell lineages than the other datasets with cell type annotations (Fig. 3a). BioID and CRAFT show reduced coverage of deeper CL terms, indicating that their annotations place a greater emphasis on less granular cell types: the mean depth of CL terms in CellLink is 5.90 versus 5.23 for BioID and 4.78 for CRAFT, in comparison to the mean depth for CL terms of 6.25 (Fig. 3b). Across all ontology depths, CellLink covers substantially more CL terms than BioID or CRAFT, and more closely matches the overall depth distribution of CL (sum of absolute differences between the normalized depth distributions = 0.157 for CellLink, compared with 0.416 for BioID and 0.517 for CRAFT). Altogether, these results indicate that CellLink captures both broad and fine-grained levels of cellular specificity.

**Fig. 3.**
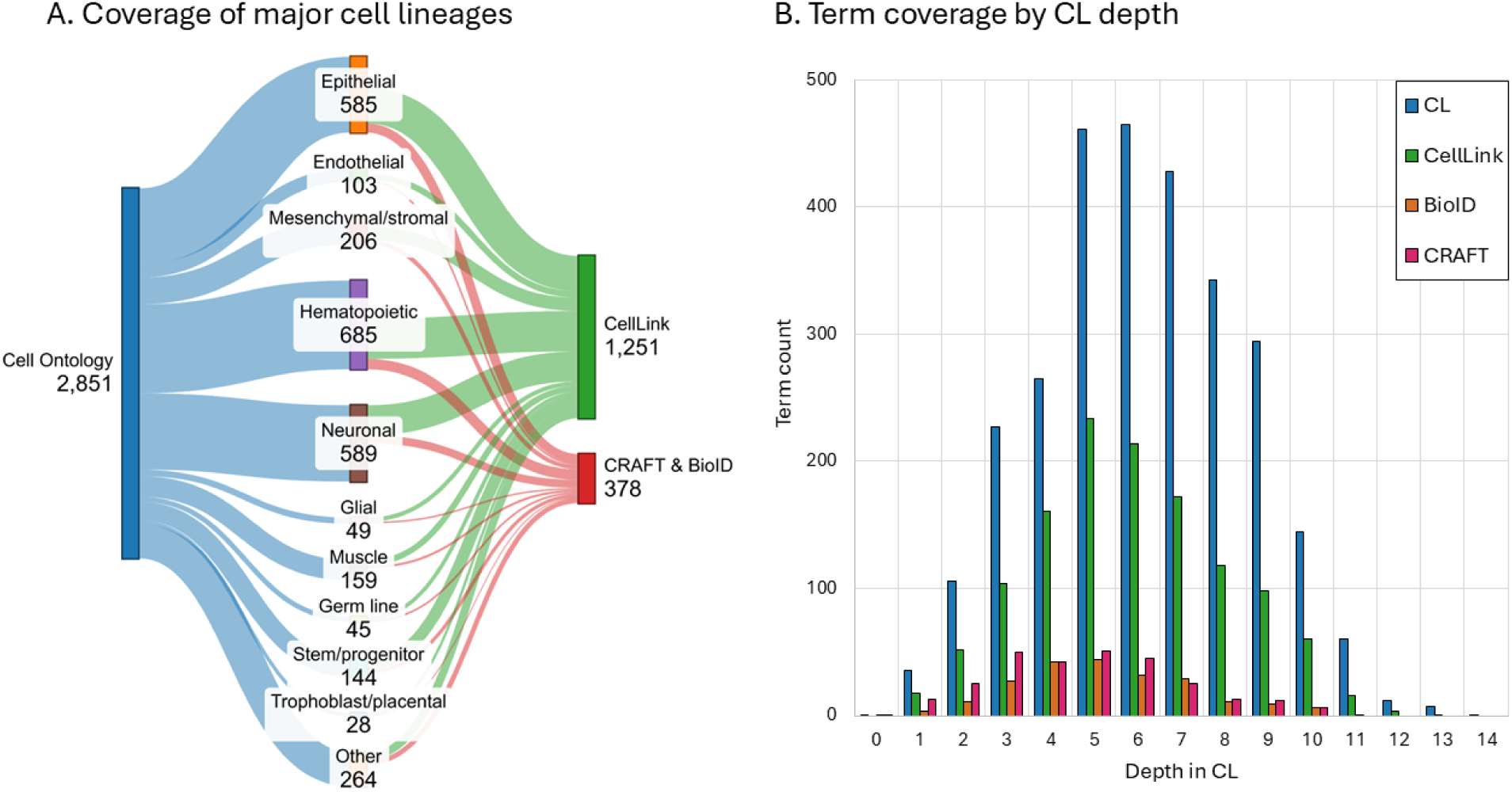
CellLink coverage of the Cell Ontology (CL). (A) Number of identifiers from the major cell lineages represented in CL, and the total CellLink and combined CRAFT and BioID datasets. (B) Comparison of the number of terms present in CL, CellLink, BioID and CRAFT at each depth in the CL ontology hierarchy. CL includes cell types from a broad range of species; cell types not present in humans or mice are included in this figure but are out of scope for CellLink.

CellLink contains a substantially higher number of low-frequency mentions than other corpora, which better reflects real-world term usage in new, emerging areas and creates a more challenging and realistic benchmark for NLP systems. Notably, CellLink contains over twice as many unique mentions that appear only once as the comparison corpora combined (Supplementary Fig. 3), with singletons comprising ∼75% of all unique mentions.

As an extrinsic validation, we conducted a five-way cross-evaluation with BiomedBERT to test whether training on CellLink generalizes across corpora. We fine-tuned BiomedBERT^7^ models on the training set of each manually annotated corpus (CellLink – combining cell phenotype and heterogeneous cell populations while dropping vague cell populations, AnatEM, BioID, CRAFT, and JNLPBA) and ran inference with each model on each of the five test sets (Table 3). As expected, each model performed best on the evaluation set for which it was trained and exhibited reduced performance on other corpora. The model trained on the CellLink corpus performed best on the evaluation set of every other corpus, excluding the models trained on the same corpus as the evaluation set, indicating its strength for training models to perform cell type NER.

**Table 3.** Cross-corpus performance of BiomedBERT NER models (F1 score)

|  |  | Training set |  |  |  |  |
| --- | --- | --- | --- | --- | --- | --- |
|  |  | CellLink | AnatEM | BioID | CRAFT | JNLPBA |
| Evaluation set | CellLink | **0.88\*** | (0.75)<sup>†</sup> | 0.52 | 0.59 | 0.66 |
|  | AnatEM | (0.69) | **0.86** | 0.49 | 0.53 | 0.47 |
|  | BioID | (0.54) | 0.36 | **0.69** | 0.50 | 0.34 |
|  | CRAFT | (0.66) | 0.58 | 0.65 | **0.82** | 0.37 |
|  | JNLPBA | (0.55) | 0.50 | 0.40 | 0.41 | **0.74** |

\*The highest score for each evaluation set is shown in bold font.
<sup>†</sup>The highest score for each evaluation set excluding the model trained on the corresponding corpus is shown in parentheses.

### Analysis of cell population naming patterns

To characterize how authors refer to cell populations, we analyzed the CellLink annotations to identify frequently used naming motifs – such as anatomical context (e.g., “peripheral”), molecular signature (e.g., “Foxp3+”), developmental stage (e.g., “immature”), or functional role (e.g., “suppressor”) – using a combination of manual curation and automated labeling (see Methods: Naming motif analysis). Because cell population names are typically hierarchical and compositional, our analysis focused on how authors combine motifs to distinguish phenotypes. We restricted our analysis to mentions linked to a single cell population, including both exact and related mentions. Therefore, vague cell populations, unlinked annotations, and annotations referring to multiple cell populations (e.g. “T and NK cells”) were excluded from this analysis.

We identified 14 motif types used as components of cell population names: root, anatomical context, lineage, molecular signature, appearance, functional role, developmental stage, state, variant, molecular signaling, disease, eponym, stimulus, and species/sex. We provide relevant examples below, see Supplementary Table 8 for complete definitions.

Cell phenotype annotations linked to a single CL identifier as an exact match showed clear lineage-dependent motif patterns compared to the baseline prevalence across all lineages combined (Supplementary Table 9). Here, prevalence refers to the proportion of mentions in which a motif type appears at least once. Cells from lineages that provide structural support – epithelial, endothelial, and mesenchymal/stromal – showed higher prevalence of anatomical context motifs (e.g., “anterior,” “liver”) and lower prevalence of molecular signature, functional role, and state motifs. Epithelial and endothelial cells also showed higher prevalence of lineage motifs, largely reflecting repeated use of the words “epithelial” and “endothelial” themselves, as these lineages lack stand-alone terms such as “neuron” or “myocyte.” Muscle cells exhibited a higher prevalence of the anatomical context motifs and a lower prevalence of the molecular signature motifs, but also had lower prevalence of the lineage motifs. Hematopoietic cells showed a profile essentially opposite of the aforementioned “structural lineages,” with high prevalence of molecular signature (e.g., “CD8+”), functional role (e.g., “regulatory”), state (e.g., “naïve”), and variant (e.g., “M1”) motifs, and significantly lower use of anatomical context motifs. The naming motif patterns for neuronal and glial cells differed substantially, despite their shared anatomical context. Neurons showed higher prevalence of the molecular signaling (e.g., “GABAergic”) and appearance (e.g., “pyramidal”) motifs and lower prevalence of lineage motifs. Glial cells exhibited very high prevalence of eponyms (e.g., “Müller”). Finally, the germ line, stem/progenitor, and trophoblast lineages showed elevated use of developmental motifs (e.g., “pluripotent”). Stem/progenitor cells also showed higher use of the lineage motif, while germ line cells showed lower use of anatomical context motifs.

Cell phenotype annotations linked as related matches, which often correspond to newly reported or incompletely characterized populations, showed clear lineage-dependent motif patterns relative to the corresponding prevalence in exactly matched cell phenotypes (Supplementary Table 10). We observed: (1) an increase in molecular signature motifs overall and across nearly all lineages; (2) overall increases in disease and state motifs; (3) decreases in the anatomical context motif in endothelial and mesenchymal/stromal cells; (4) increases in the developmental motif within the epithelial and endothelial lineages; and (5) decreases in the prevalence of the appearance motif in neuronal cells and eponyms in glial cells.

Finally, comparing the heterogeneous cell population mentions, combining both the exact and related matches (Supplementary Table 11) to the corresponding prevalence in exactly matched cell phenotypes showed a substantial increase in the overall prevalence of disease motifs (from 0.008 to 0.169 motifs per annotation) and a small overall increase in anatomical context motifs. We also observed an overall decrease in the prevalence of molecular signature motifs, particularly within the hematopoietic lineage.

### Evaluation of NLP models using CellLink

The utility of the CellLink corpus to benchmark NLP systems for NER and entity linking (EL) to the CL reference was assessed. We evaluated three systems for NER on our dataset: a BiomedBERT-based system using three separately fine-tuned models (one per entity type, with predictions combined) and two GPT models (GPT-4.1^20^ and GPT-5.2^21^), prompted to simultaneously recognize all three types from examples. BiomedBERT achieved high performance for cell phenotypes, with a 0.874 F1 score for exact spans, but had more difficulty with heterogeneous and vague cell populations (Table 4). GPT-4.1 and GPT-5.2 both performed moderately well overall with approximate spans and merged labels (0.791 and 0.788 F1 score, respectively) but had trouble identifying the exact spans (0.634 and 0.656 F1 score, respectively) and distinguishing between entity types. The performance differences between GPT-4.1 and GPT-5.2 were small and inconsistent across settings. We anticipate that additional prompt engineering, especially adjusting the number of examples provided in the prompt (few-shot learning), or fine-tuning would improve the results of the GPT model for NER.

**Table 4.** Performance of NER models on the CellLink test set (exact F1 score* / approx. F1 score^†^)

|  | BiomedBERT, fine-tuned | GPT-4.1, few-shot | GPT-5.2, few-shot |
| --- | --- | --- | --- |
| Cell phenotype | 0.874 / 0.941 | 0.612 / 0.738 | 0.568 / 0.768 |
| Heterogeneous cell population | 0.703 / 0.779 | 0.240 / 0.351 | 0.257 / 0.419 |
| Vague cell population | 0.592 / 0.696 | 0.230 / 0.305 | 0.163 / 0.240 |
| All | 0.834 / 0.906 | 0.509 / 0.627 | 0.477 / 0.571 |
| Merged labels<sup>‡</sup> | 0.863 / 0.926 | 0.634 / 0.791 | 0.656 / 0.788 |

\*Exact F1 score refers to a match in which the predicted span and entity type exactly match the annotated span and entity type.
<sup>†</sup>Approx. F1 score refers to a match in which the predicted entity type exactly matches the annotated entity type, and the predicted span overlaps the annotated span. Each predicted annotation may match at most one reference annotation, and vice versa.
<sup>‡</sup>Merged labels refers to combining all annotations into a single type.

For EL, we evaluated three embeddings-based models (SapBERT^22^, MedCPT-Query-Encoder^9^, and OpenAI’s text-embedding-3-large^23^) and an agent-based linker (using GPT-5.2^21^ as the underlying language model), on the ground truth mentions of the test set for EL to CL (cell phenotype and heterogeneous cell population annotations only) (Table 5). The GPT agent achieved higher performance than the embeddings-based models for all evaluations, except for linking heterogeneous cell populations when an exact match was available. Overall, the agent achieved an F1 score of 0.782, compared with SapBERT, the next-highest performing model, at 0.718. As expected, all models performed better when linking to an annotation that has a single, exact-match identifier than annotations linked to multiple identifiers. Performance of all models dropped dramatically for heterogeneous cell populations when allowing annotations that have multiple identifiers, possibly because heterogeneous cell population annotations are sometimes linked to many related identifiers. All models performed better on exact match EL for heterogeneous cell populations than for cell phenotypes (0.072 increase in F1 score, on average), possibly because heterogeneous cell population annotations are less likely to be context-dependent than cell phenotype annotations.

**Table 5.** Evaluation of multiple entity linking models on the CellLink test set (F1 score)

|  |  | SapBERT | MedCPT-Query | OpenAI text-embedding-3-large | GPT-5.2 Agent |
| --- | --- | --- | --- | --- | --- |
| Exact matches only | Cell phenotype | 0.836 | 0.820 | 0.806 | **0.891\*** |
|  | Heterogeneous cell population | **0.956** | 0.846 | 0.931 | 0.909 |
|  | Overall | 0.842 | 0.821 | 0.812 | **0.892** |
| All IDs | Cell phenotype | 0.738 | 0.735 | 0.718 | **0.805** |
|  | Heterogeneous cell population | 0.520 | 0.506 | 0.510 | **0.559** |
|  | Overall | 0.718 | 0.714 | 0.699 | **0.782** |

\*The highest score for each evaluation is shown in bold.

While the GPT-5.2 agent outperformed the other models, it was significantly slower and more expensive to run. SapBERT and the MedCPT-Query-Encoder ran locally on a p100 GPU in 1.47 and 1.30 minutes, respectively. OpenAI text-3-embedding-large ran in 3.68 minutes, limited by Microsoft Azure’s processing rate quota. The GPT-5.2 agent took over two hours to run, limited by a Microsoft Azure quota and calls to the Ontology Lookup Service^24^.

We also evaluated the performance of embedding-based models – SapBERT, MedCPT, and text-embedding-3-large – for EL using top-k recall: the recall at the k highest-scoring results. The agent was not evaluated in this paradigm because it is designed to only return a single identifier. This evaluation used the ground truth mentions and considered cell phenotype and heterogeneous cell populations without differentiating between the two, nor between exact and related identifier matches. SapBERT again achieved the highest performance for the top-1 identifier. However, the difference between the three models for top-5 is negligible (0.003), and OpenAI’s text-embedding-3-large achieved the highest performance for top-10 (Supplementary Table 12).

We analyzed the relationship between the confidence of SapBERT in its top-1 identifier and whether an annotation was novel/unreported (i.e., the annotation was not linked to a CL identifier with an exact match). SapBERT had high confidence (>0.95) in the vast majority of exact-match identifiers, whereas its confidence for linking annotations with related-match identifiers or no IDs was mostly between 0.40−0.85 (Fig. 4a). SapBERT confidence had an area under the receiver operating characteristic curve (AUROC) of 0.867 for determining whether a mention was mapped to an ontology term, which was similar for both cell phenotype and heterogeneous cell populations (0.857 and 0.894, respectively; Supplementary Table 13). Additionally, SapBERT showed high capacity to determine whether it was correctly matching an identifier from the reference vocabulary (Fig. 4b, AUROC=0.89), which indicates that thresholding SapBERT confidence could be useful for simultaneously identifying novel cell types and increasing the precision of a cell type text entity linking system.

**Fig. 4.**
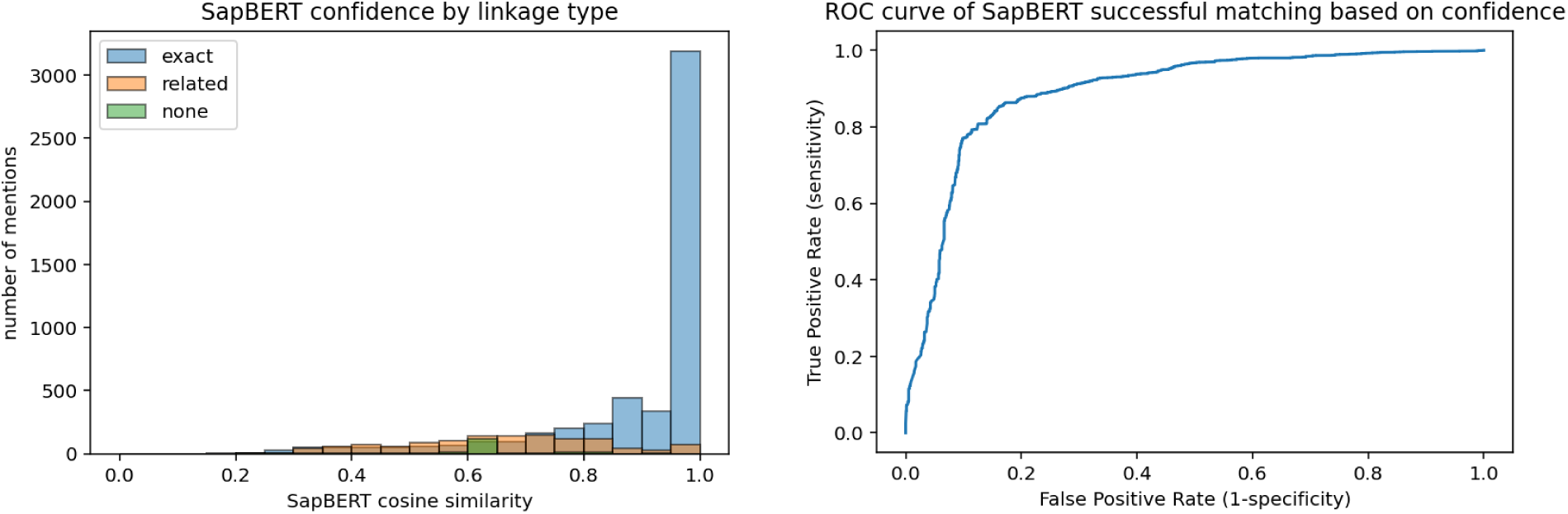
SapBERT cosine similarity scores reflect type of EL match and prediction accuracy. (A) Distribution of top-ranked SapBERT cosine similarity scores across annotations with an exact match, related matches, or no matches (no identifier assigned). (B) Receiver operating characteristic (ROC) curve quantifying the ability of the top-ranked SapBERT similarity score to distinguish correctly predicted exact matches, i.e., annotations corresponding to ontology terms. Area under the ROC curve = 0.893.

### Use of CellLink to expand the Cell Ontology

CL aims for comprehensive coverage of well-characterised, non-pathological, *in vivo* cell types, with high coverage of mammalian cell types^25^. The CellLink corpus, particularly the cell phenotype annotations without an exact match to a CL record, provides an opportunity to identify candidate cell types from the literature that may warrant inclusion or revision in CL.

Despite its focus on populations of natural cells, the CellLink corpus includes cell population mentions spanning a wide range of experimental and disease contexts. To align with the CL scope, cell names were filtered using keywords indicating artificial or abnormal cell types or states (such as “cultured”, “iPSC-derived”, “abnormal”, “infected”). From the remaining annotations, we reviewed phenotypes lacking exact matches to CL terms. For example, this process identified several cell phenotype annotations in CellLink with related matches to the general term chondrocyte that were not yet represented in CL, indicating a gap in coverage.

When incorporating transcriptomically-defined cell types, CL curation requires phenotypes to map back to classical cell types, demonstrate phenotypic consensus across multiple independent studies, or be characterized using complementary modalities such as spatial localization or morphology. Applying these criteria, the chondrocyte phenotypes identified in the CellLink corpus, together with their textual context, were used to support a targeted expansion of the chondrocyte branch of CL, aligning it with evidence from modern single-cell, lineage tracing, and spatial transcriptomic data. This effort involved restructuring part of the hierarchy and adding missing high-level classes and functional subtypes, such as “regulatory chondrocyte,” “homeostatic chondrocyte,” and “effector chondrocyte,” originally identified in single-cell RNA-sequencing atlases of human cartilage ^26^ and subsequently validated across independent datasets (Fig. 5).

**Fig. 5.**
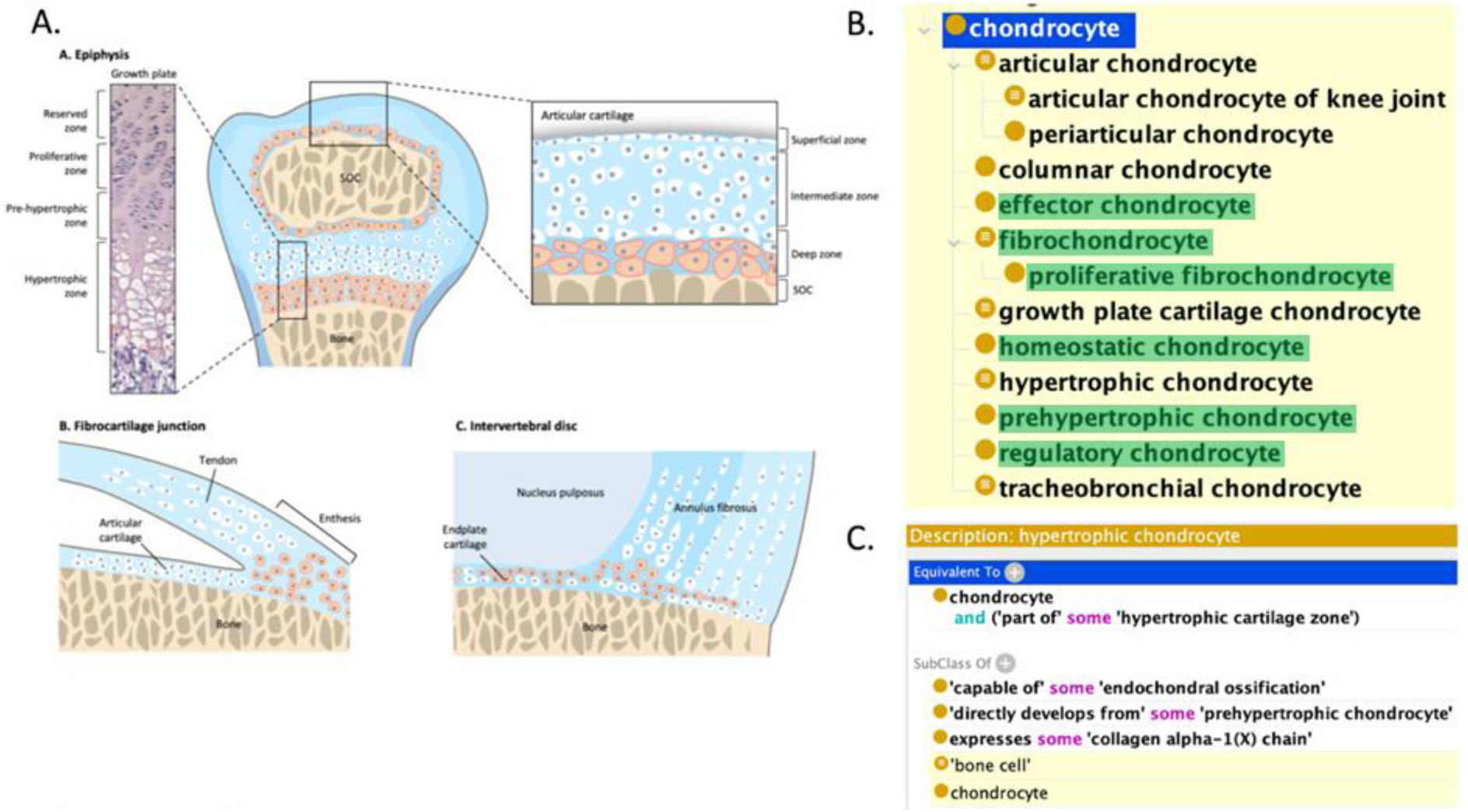
Expansion of the chondrocyte branch in the Cell Ontology (CL) using NLM CellLink. (A) A schematic of chondrocytes across cartilage regions, adapted from Chen et al.^27^ under the Creative Commons Attribution 4.0 International License (https://creativecommons.org/licenses/by/4.0/). (B) An expanded and restructured hierarchy of chondrocytes in CL, including newly added terms highlighted in green (prehypertrophic chondrocyte, regulatory chondrocyte, homeostatic chondrocyte, effector chondrocyte, fibrochondrocyte, proliferative fibrochondrocyte, fibrochondrocyte progenitor cell*, osteochondral skeletal stem cell*, and hybrid osteochondral skeletal cell*) and revised existing terms. (C) Logical definition of the term “hypertrophic chondrocyte”, which was revised based on CellLink passage information. *These cells were added into CL but are not displayed in the hierarchy as they represent developmental links.

In addition to identifying missing classes, we also revised outdated definitions for existing CL terms. For example, “hypertrophic chondrocyte,” historically defined as a terminally differentiated cell, was updated to reflect lineage-tracing and trajectory evidence demonstrating that these cells can transdifferentiate and contribute to osteogenic progenitor lineages^28^. Conversely, certain candidates were excluded due to insufficient consensus or ambiguous usage in the literature. We excluded “dedifferentiated chondrocyte,” as this term is used to refer to both a reversible *in vitro* phenotype associated with monolayer culture^29, 30^ and a transient biological state during skeletal stem/progenitor cell (SSPC) development^28^. Similarly, “chondrocyte-like osteoprogenitors”^31^ were not included at this time because current evidence remains limited and the nomenclature is used inconsistently across studies^31, 32^, precluding representation as a distinct CL term.

CL adds tissue-specific cell type terms wherever possible, but the anatomical coverage of cartilage regions and types in spatial transcriptomics studies is currently insufficient to support adding these assertions to all but two of the added types^33, 34^. As spatial atlases expand, we intend to revisit this branch in CL to add more tissue-specific subtypes. This chondrocyte case study demonstrates how CellLink can support scalable, domain-focused ontology refinement by supporting identification, evaluation, and curation of cell phenotypes from the literature, helping the ontology to remain aligned with the increasing granularity of modern single-cell studies.

## Discussion

The rapid adoption of single-cell technologies has significantly shifted how cell populations are referenced in the literature, necessitating resources that reflect the increasing complexity of cell nomenclature. In this work, we present CellLink, a large-scale corpus of human and mouse cell population mentions mapped to CL. Unlike prior resources, CellLink distinguishes between three types of cell populations — specific cell phenotypes, heterogeneous cell populations, and vague cell populations — and links annotations to CL in a context-dependent manner with explicit exact and related qualifiers. We validated corpus quality through inter-annotator agreement and assessed utility on NER and EL tasks and on CL expansion. By systematically capturing the variety and granularity of cell phenotypes in the modern literature, this resource reflects the complexity of cell nomenclature more comprehensively than any previous corpus. As timely scientific infrastructure, CellLink provides a foundation for automated extraction of cell-centric knowledge from the literature, thereby facilitating ontology expansion and the construction of knowledge resources connecting cell phenotypes to genes, diseases, and anatomical structures in the future.

Our work demonstrates that the cell populations referenced in the literature do not necessarily correspond to discrete biological entities, but instead reflect the information available to authors, the conventions of their field, and the methods used to generate the data. Many of the elements that appear in cell type names can be ambiguous: anatomical context may or may not indicate a distinct phenotype; disease qualifiers are used inconsistently and the impact of disease context on cell phenotypes is often unclear; numerical labels may refer either to well-established subtypes or to arbitrary cluster identifiers; and marker-based names arising from scRNA-seq often describe transcriptionally defined clusters that may not represent stable cell types. For example, disease-associated cell phenotypes sometimes reflect reversible changes in normal cells that occur in response to the presence of a disease process, but in other cases they reflect stable cellular abnormalities that contribute to the physical basis of disease; this distinction is often not apparent in the name. Moreover, because scRNA-seq workflows typically annotate clusters using marker genes and may split known phenotypes into multiple subclusters, authors frequently use combinations of marker genes to distinguish closely related populations, even when it is unclear whether these distinctions are biologically meaningful. As a result, interpreting a cell population name often requires substantial contextual knowledge, and naming alone is not a reliable indicator of whether the authors intended to denote a new cell phenotype, a transient state, or simply a cluster-level distinction.

Creating CellLink required numerous judgment calls that highlight the tension between reproducibility and biological nuance. Even with detailed guidelines, annotating cell type mentions inevitably involves interpreting biological context and author intent. The annotators reported that disagreements often reflected alternative, equally reasonable readings rather than clear errors. The moderate initial inter-annotator agreement (0.69) thus underscores the intrinsic ambiguity of both language and biology rather than inconsistency in the process. Through iterative discussion and refinement, all differences were reconciled, producing a consensus that balances principled rules with practical flexibility. This balance – structured guidance complemented by expert discretion – is essential for achieving both consistency and biological accuracy in biomedical corpora.

Our analysis of the naming motifs in cell population mentions demonstrated that authors draw on a diverse set of attributes when referring to cell phenotypes, and that these naming conventions vary across biological lineages. For cell phenotypes that link exactly to CL terms, these patterns largely reflect historical and methodological practices across the disciplines that study each lineage. Epithelial, endothelial, mesenchymal/stromal, and muscle cells primarily perform structural functions tied to their anatomical location, and were historically characterized through histological staining^35^. The historical controversy over the developmental origin of endothelial cells likely contributes to their increased use of lineage motifs^36^. In contrast, hematopoietic cells – including immune cells – are highly migratory and undergo multiple differentiation events. Their identities are therefore more closely associated with functional roles (e.g., “helper,” “cytotoxic”), molecular signatures (particularly the “cluster of differentiation” CD system), cell states, and variants (e.g. “T helper 1”)^37^. Neuronal names frequently incorporate morphology (appearance motifs) and specific neurotransmission properties (molecular signaling motifs), reflecting early characterization with microscopy and later by functional signaling properties^38^. Glial cells show unusually high rates of eponyms, likely a legacy of 19^th^ century naming practices and their historically unclear function^38^. Finally, cells from the germ line, stem/progenitor, and trophoblast lineages are defined by their role in reproduction and their position along differentiation trajectories, explaining the higher use of developmental-stage motifs^39^. Stem/progenitor cells also exhibit higher rates of lineage motifs, which identify the specific developmental trajectory.

The shift in motif usage in cell phenotypes between exact and related matches suggests that many related matches correspond to phenotypes identified using recent experimental technologies. Increased emphasis on molecular signatures, state descriptors, and disease context is consistent with high-resolution single-cell transcriptomic profiling. Related matches also show reduced reliance on anatomical context in endothelial and mesenchymal/stromal cells, reflecting the emergence of transcriptionally defined subtypes across tissues. Conversely, epithelial and endothelial related matches show greater use of developmental motifs, consistent with single-cell studies capturing intermediate differentiation states. For neuronal and glial phenotypes, related matches exhibit fewer morphology-based and eponym-based terms, respectively, paralleling the shift from classical, morphology-driven nomenclature toward molecular and functional classification. CellLink therefore captures terminology associated with both classical phenotypes and emerging fine-grained / transitional populations.

The substantially increased use of disease motifs in heterogeneous cell population names is largely dominated by cancer cells, a common subject of scRNA-seq studies. The increase in anatomical context terms is consistent with names that describe mixed populations in the tissue of origin. Conversely, the reduction in molecular signature motifs aligns with the fact that these mixed populations would not be expected to show shared marker profiles. Together, these differences underscore that heterogeneous cell populations function as context-dependent groupings of convenience rather than discrete cell types with shared phenotypic characteristics, supporting their treatment as a separate category.

Overall, the analysis of naming motifs demonstrates that the naming of cell populations is increasingly descriptive and compositional, especially in studies using high-dimensional profiling technologies. Representing additional perturbations, such as disease context, environmental response, or experimental modification, would further extend this trend. As compositionality increases, it becomes impractical to enumerate all possible combinations as pre-coordinated ontology terms, suggesting that future cell population representations may benefit from approaches that support composing descriptors logically.

Several characteristics make the CellLink corpus a valuable benchmark for evaluating the robustness of biomedical NLP and natural language understanding systems. Our passage selection process yielded a high proportion of unique mentions and identifiers; the 1,251 unique CL identifiers annotated (Table 1) represent approximately 44% of the concepts in CL v2025-01-08. This variety should support rigorous method evaluation and promote model generalization. The corpus includes three entity types that are semantically distinct but have high surface-level linguistic similarity, making lexical cues alone often insufficient for correct classification. Distinguishing between cell phenotypes and heterogeneous cell populations will likely require models to have strong prior or external knowledge. Vague cell population annotations are more like entity descriptions rather than traditional, concrete named entities. Recognizing vague cell population annotations and annotations where the authors omitted essential words such as “cell” (or other root words) will likely require models to capture deeper contextual and semantic cues. The annotated CL identifiers were linked using the meaning of the mention in context, so that the mention “fibroblasts” in a context where it clearly refers to “cardiac fibroblasts” (e.g., PMID:31222588) was linked to the CL term fibroblast of cardiac tissue (CL:0002548). This context-dependent linking will substantially limit the effectiveness of rule-based and dictionary-driven approaches, as lexically identical mentions may be linked to different CL terms in our corpus. Moreover, CellLink annotations include both exact and related matches to CL terms, requiring systems to not only retrieve terms but also assess the degree of the match. Finally, the inherently compositional structure of cell type names means that resolving the individual mentions within coordination ellipses – compressed phrases referring to multiple related entities, such as “hematopoietic stem and progenitor cells” and “CD4^+^ and CD8^+^ T cells” – provide a natural test of whether systems can decompose and ground complex, hierarchical mentions.

This work has several limitations and natural next steps. The corpus focuses on human and mouse literature published between 2019 and 2024 and prioritizes naturally occurring cells over experimentally modified cells. Annotation boundaries for descriptive phrases necessarily involved judgment, particularly for vague populations, and links to CL depend on the ontology version available at the time of annotation. Our evaluation primarily used out-of-the-box NER and EL tools, but more advanced methods will be needed to take full advantage of the dataset. For NER, it will be necessary to handle overlapping mentions, resolve coordination ellipses, and utilize context cues to interpret vague cell populations. For EL, future work includes fine-tuning linking models, developing context-dependent systems, building classifiers to distinguish exact from related or absent matches, and determining automatically how many related identifiers should be associated with a given annotation. Our analysis provides preliminary support for these efforts by demonstrating that SapBERT confidence serves as a strong preliminary predictor of the accuracy of exact-match linking.

Given the combination of consistent annotation and expert judgement, CellLink provides a valuable resource for developing automated systems to identify mentions of human and mouse cells in the biomedical literature. By distinguishing multiple types of cell populations and linking them to CL identifiers through both exact and related matches, CellLink facilitates the integration of literature-mined data and structured biological knowledge. This approach supports biocuration through novel cell type and synonym discovery, thereby contributing to the continued development of CL. CellLink could also be applied for PubMed information retrieval systems, such as PubTator^8^, enabling users to perform more efficient, comprehensive searches for specific cell types. CellLink provides critical groundwork for relation extraction and knowledge graph construction, connecting cell phenotypes with other important biological entities including genes, mutations, anatomical structures, and diseases.

## Methods

To maximize the value of expert biological annotation, we automated as much of the preparatory work as possible prior to annotation. Automated sampling and pre-annotation ensured a focused selection of text enriched in relevant cell populations, allowing domain experts to focus their efforts on the more challenging task of interpreting the cell populations described in the selected text. The overall workflow included (1) article retrieval and passage selection, (2) expert annotation following an iterative annotation process, (3) shared guidelines to ensure quality, and (4) benchmarking and use of the resulting corpus.

### Article retrieval and passage selection

Our prior work demonstrated that full-text articles contain substantially more specific content than titles and abstracts alone^19, 40^. However, annotating entire articles requires significantly more expert time and effort. Individual sentences, on the other hand, often lack sufficient context for correct annotation. Passages, which typically correspond to a paragraph, provide more context than sentences and much greater variety per unit of effort than full articles, especially if passages are selected judiciously. We therefore chose passages as the annotation unit as a practical middle ground.

We created a document pool using a PubMed query designed to retrieve recent studies using single-cell genomic technologies, especially transcriptomics, as well as broader discussions of cell types (Fig. 1):

(“single cell*” OR “scRNA*” OR “single nucleus” OR “snRNA*” OR “single nuclei” OR “cell type*” OR “spatial transcript*“) AND (human OR mouse) AND (“2019“[Date - Publication] : “2024“[Date - Publication]) NOT (“preprint” [Publication Type])

The query limits the results to articles published in 2019 through 2024, inclusive, to reflect the rapid growth of single-cell methods during this period. We focused on human and mouse studies due to their central role in biomedical research. The search, conducted via eUtils^41^ on Dec 31, 2024, returned 70,859 PubMed identifiers (PMIDs); 35 books were filtered, leaving 70,824 articles. Articles matching these PMIDs in the PubMed Central (PMC) Open Access subset with CC-BY or CC0 licenses were retrieved as full text (n=40,793); for all other articles, only the titles and abstracts were retrieved (n=30,031). Retrieval was performed via the PubTator API^8^.

Passages, as defined by the PubTator API, include individual paragraphs, section headers, captions, and tables. After retrieval, we filtered out passage types that were not found to contain relevant cell type mentions: methods, references, funding, acknowledgements, author contributions, competing interests, and review information. Extremely long tables were also filtered. About 1.5 million passages remained after filtering. Of the 4.9 million passages initially retrieved, approximately 40% were references. Retained passages included titles, abstracts, introductions, results, discussions, conclusions, figure captions, supplementary information, abbreviations, and tables.

To ensure broad topical coverage and balance common versus less frequent subjects, we grouped articles into clusters using MeSH terms from the Anatomy (A), Organisms (B), and Diseases (C) branches of the MeSH hierarchy. Clusters represented anatomical systems (e.g., “Nervous-Anatomy”) or diseases of a specific system (e.g., “Cardiovascular-Disease”). Clusters were defined by a small set of MeSH terms, with all descendants included (Supplementary Table 1). Articles may belong to multiple MeSH clusters, and since MeSH terms are assigned to articles rather than specific passages, the content of each passage may not perfectly match its cluster. These clusters were subsequently included as features informing passage selection to promote broad and balanced coverage across biomedical topics.

Passages were selected using a sampling strategy that balanced representativeness with enriching less-frequent values. For each passage, we extracted a set of discrete features derived from article-level metadata (journal, year, MeSH terms, and MeSH clusters), passages attributes (type, length, and tokens present), and semantic annotations (entities from PubTator and cell type predictions from BiomedBERT models trained on each comparison corpus). For each feature type, a target distribution was initially estimated from the pool of filtered passages, then adjusted to reflect annotation priorities. Specifically, we emphasized long-tail values by applying temperature scaling (T=1.6)^42^, increased the number of natural human and mouse cells (e.g., increasing the target frequency of passages containing cell population predictions by 800%), and reduced selection of out-of-scope content, including passages focused on cancer (e.g., by reducing the frequency of the “Cancer-Disease” MeSH cluster by 50%) and cell lines (e.g., by reducing the frequency of PubTator cell line annotations by 50%). We also increased the average passage length by 300%. The complete set of target distribution adjustments is provided in Code availability. Passages were then greedily selected to minimize the Kullback-Leibler (KL) divergence^43^ between the feature distribution of the selected set and the adjusted target distribution, with a maximum of two passages selected per article. Selected passages were manually screened and excluded only if they contained irrelevant content (e.g., non-target species; approximately 1%). Pseudocode for the passage-selection procedure is provided in Supplementary Algorithm 1.

### Annotation process

Four experienced cell biologists annotated passages following a shared set of annotation guidelines, summarized below. Each passage was pre-annotated using a BiomedBERT^7^ model fine-tuned on AnatEM^14^ for NER and SapBERT^22^ for zero-shot EL to terms and identifiers in CL. The pre-annotated spans and identifiers were provided to the annotators as candidate annotations, but annotators exercised full control over the final annotations. The annotation system required annotators to explicitly accept or correct pre-annotations they wished to retain. Annotators read each assigned passage in full, ensuring cell populations missed by the pre-annotation were captured. Annotation was performed in TeamTat^44^, using the EBI Ontology Lookup Service (OLS)^24^ to search CL.

Annotation training consisted of repeated rounds of independent annotation, followed by collaborative resolution, allowing annotators to refine the guidelines and adjust their annotations for consistency. Following the training phase, each passage was annotated by two annotators in a 3-round annotation process, as described in our prior work^19^:

1. Independent annotation: annotators labeled passages without seeing the other’s annotations.
2. Independent resolution: annotators reviewed their partner’s annotations and revised their own annotations, without discussion or knowing their partner’s identity.
3. Collaborative resolution: Annotators met to discuss and resolve any remaining differences.

This balances independent work, which reduces bias, with collaborative discussion to enable consistency and ensure consensus. Passages were separated into daily batches of equal size that paired each annotator with every other annotator.

After all annotation batches were completed, potential inconsistencies were flagged for re-review. Two types of potential inconsistencies were automatically identified. The first involved mention-identifier pairs that diverged from the predominant annotation pattern for the mention, suggesting a possible incorrect identifier. The second involved phrases containing an annotated cell population (e.g. “epithelial cell”) followed by a subset descriptor (such as “subsets”) where the entire phrase should be annotated as a vague cell population. Flagged cases were re-reviewed to ensure consistency.

IAA was calculated as the F1 score between the annotations created by the two annotators assigned to each passage, based solely on the first-round annotations, to capture baseline agreement before discussion. Several variants were computed, defining agreement in different ways:

- Strict span-identifier agreement: Exact match in annotation span, entity type, and linked identifier.
- Strict span agreement: Ignores the identifier.
- Strict identifier agreement: Presence of the linked identifier anywhere within a passage.
- Approximate span agreement: Exact entity type and any overlap between the annotations.

For IAA, related-match identifiers were treated as “no ID” because there are usually several acceptable combinations of related identifiers.

### Overview of annotation guidelines

The annotators followed guidelines summarized in Methods: Annotation process and described in full in Supplementary File 1. In short, they identified all mentions of cell populations and selected the shortest span that fully captured the distinct population. Descriptors relevant to cell identity, such as state (e.g., proliferating), marker genes (e.g., CD4), and anatomical locations (e.g., lung), were included when present.

Three entity types were distinguished:

- Cell phenotypes: Unambiguous mentions of distinct, stable cell types and their transient states (if specified). Examples: “fibroblast”, “activated fibroblast”.
- Heterogeneous cell populations: Groups of cells that share some characteristic but are from multiple distinct developmental lineages. Examples: “immune cells”, “kidney cells”.
- Vague cell populations: Mentions of cell populations that cannot be identified precisely. Example: “Ly6a, Cebpb, and Egfr expressing cells”, “microglia-like cells”.

Each annotation was classified as one of these three entity types. Cell lines were excluded to focus on naturally occurring cells.

Annotations classified as cell phenotypes or heterogeneous cell populations were linked to identifiers in CL (version 2025-01-08). Annotations classified as vague cell populations were not linked, since their identities cannot be determined precisely. Annotators searched CL for a term with the same meaning as the annotation; these matches were tagged as exact. If no such term was available, one or more related matches could be linked, including terms that were broader, narrower or otherwise related, based on annotator judgement. If no CL terms were judged to be closely related, then no identifier was assigned.

In the literature, authors often omit details that can be inferred from context. For instance, “lens fiber cells” may be shortened to “fiber cells” when the lens context is clear and unlikely to be confused with other cell types commonly described as fibers, such as myocytes. Annotators therefore considered the entire passage – and, if necessary, the entire article – to determine the most appropriate identifier. An article that referred to “CD56^−^ NK cells” (note the superscript minus) and later referenced the same population as “CD56^−^ cells,” for example, would have both mentions linked to the same identifier. Similarly, “inhibitory neurons” in humans would be linked to the CL term for “inhibitory interneurons” in most contexts. Consequently, the same mention text could be assigned different identifiers, depending on the context.

The annotators recorded questionable or potentially inaccurate cell populations as the authors appeared to intend, without correction. For example, all non-redundant descriptors were retained in the span, including anatomical locations, even when their biological relevance was debatable (e.g., whether T cells differ by location). This approach kept the task focused on interpretation of author intent rather than correction. Special cases were handled as follows:

- Coordination ellipses: If part of a term was omitted through conjunction (e.g., “immune and endothelial cells”), the entire span was annotated as a single annotation and linked to the identifiers for each type, separated by semicolons.
- Cell ellipses: If “cell” or another root term is omitted but implied because the context clearly indicated a cell population, the span was annotated. For example, “epithelial markers” implies “epithelial <u>cell</u> markers,” so the word “epithelial” would be annotated. These cases required annotator judgment, since terms like “epithelial” can refer to other entities, such as tissues.
- Subset descriptors: Terms such as “subset,” “subtype,” “subpopulation,” or “class” imply an unspecified subset of a base cell type (e.g., “neuron subtypes”) and were typically annotated as vague cell populations.
- Overlapping spans: Overlapping annotations were allowed only when a vague cell population contained a nested cell phenotype or heterogeneous population annotation (e.g., “neuron” inside “neuron subtypes”). The corpus thus remains usable if vague cell population annotations are filtered.
- Cell mentions within larger phrases: Cell populations mentioned within noun phrases that were out-of-scope were still annotated to capture the cell concept. For example, the mention “ovarian cell” in “Chinese hamster ovarian cell line” was annotated as a heterogeneous population.

### Annotation tradeoffs and scope decisions

Annotating cell population mentions required judgments balancing principles and pragmatic constraints to maintain both consistency and biological accuracy. Each annotation in CellLink linked to a CL term is labeled as exact or related, making one of the more significant judgments determining the degree of relatedness between entities. While every cell can exhibit some phenotypic variation due to environmental context, overuse of the related qualifier would dilute its meaning. For example, in an article discussing cells from patients with diabetic retinopathy, vascular endothelial cells and neurons may express altered phenotypes in this diseased context; whether these cells should be considered exact matches is debatable. Similarly, annotators had to judge whether a cell population mention described a vague cell population, particularly when authors used qualifiers such as “-like” in cases where the available evidence was insufficient to authoritatively characterize the cell type. For example, the annotators labeled the mention “Th1-like IL-17+ cells” (PMID: 37215131) as a vague cell population. However, many cases were more straightforward. For example, overly general mentions such as “lung cells” were not linked to narrower terms like “lung epithelial cell,” and “immune cells” were not linked to “leukocyte,” which represents only one lineage of immune cells.

Additional judgment was required to determine which descriptors to include within the annotation span. Mouse strains and cell lines, such as “HEPG2” in “HEPG2 <u>liver cancer cells</u>,” were excluded as out of scope. However, we included descriptors indicating an organ location (e.g., the entire phrase of “<u>liver</u> <u>memory T cell</u>”) and association with tumors (e.g., the entire phrase of “<u>tumor infiltrating lymphocytes</u>”). While it is often unclear whether these descriptors modify the cell phenotype – for example, whether the phenotype of liver memory T cells is distinct from memory T cells in other tissues – we retained them to remain faithful to the meaning intended by the author.

### Corpus usability evaluation

We fine-tuned and applied various models for multiple NER and EL tasks. The corpus was divided into 1/2 training, 1/6 validation/development, and 1/3 testing. This division was performed using an iterative hill-climbing approach, with the intention of ensuring that MeSH clusters and annotations from each entity type were proportionally divided, and to reduce the overlap between the training and test sets by increasing the number of mentions and identifiers unique to each set. Formal definitions of the evaluation metrics are provided in Supplementary Note 1 and source code for all models compared is provided as described in Code availability, including the prompts used for large language models.

We fine-tuned BiomedBERT models independently on each of the following corpora: CellLink, AnatEM, BioID, CRAFT, and JNLPBA. Before fine-tuning, we filtered the non-cell types from each corpus. In the CellLink corpus, we merged cell phenotype and heterogeneous cell population annotations and removed vague cell population annotations to attempt to match the annotation styles of the other corpora.

Independently, we fine-tuned a BiomedBERT model for each of the three entity types in CellLink: cell phenotype, heterogeneous cell population, and vague cell population. Each model was fine-tuned on the training set, using the annotations of the corresponding type, and applied to the testing set. Their output was pooled to give a single set of predictions for each document. We also applied GPT-5.2, prompting the model to simultaneously identify all three entity types. Examples of each entity type were provided, but without the surrounding passage for context. OpenAI’s structured outputs functionality was used to ensure a consistent format.

EL was performed using three embeddings-based models – SapBERT, MedCPT-Query-Encoder, and OpenAI’s text-embedding-3-large – and an agentic AI approach we developed using the pydantic_ai Python library^45^, modeled after prior work demonstrating agentic approaches^46^. This agent, powered by GPT-5.2, queries CL via OLS^24^ through the Ontology Access Kit^47^. All evaluations used the gold-annotated mentions, which were preprocessed to expand abbreviations identified in the full article using Ab3P^48^ and to apply a conservative plural stemmer^49^.

Under the single-link EL paradigm, each mention was embedded, and the CL term or synonym (previously embedded using the same model) with the highest cosine similarity was selected as the linked identifier. For top-k linking, each mention was linked to the *k* CL terms whose embeddings had the greatest cosine similarities to the mention embedding. The EL agent was instructed to search CL for a term that matched the given annotation and to search for synonyms or related terms if no exact match is found. Example search queries made by the agent are available in Supplementary Table 14. The agentic approach used the same abbreviation expansion and plurals stemming used for the embedding models.

EL was performed for each model using the ground truth linkages in the test set, limited to the cell phenotype and heterogeneous cell population annotations (since vague cell population annotations are not linked). We evaluated the performance of each model on each entity type and by the top-1, top-5, and top-10 ranked results (Supplementary Table 12). The top-k evaluations excluded the AI agent, which always returns one identifier. We used F-score as the metric for the single-link evaluation and recall as the metric for the top-k evaluation. We also evaluated the performance of each model after filtering annotations that did not have an exact match. A 100% performance is unattainable without this filtering under the single-link paradigm where each model links exactly one identifier per annotation because some mentions have multiple related matches.

Lastly, we analyzed ROC curves and AUROC to assess the ability of the cosine similarity score of the top-1 result using SapBERT embeddings to (1) discriminate between exact, related, and unlinked annotations, and (2) as a threshold to tradeoff between precision and recall in the top-1 identifier.

### Naming motif analysis

The individual components of cell population names, which we termed naming motifs, were identified for CellLink annotations and for names in CL using an iterative, semi-automated workflow. CellLink annotations that were not linked to a CL term, including all vague cell-population mentions, and spans referring to multiple concepts (e.g., “T and NK cells”) were removed, leaving 18,633 (83.3%) of the original 22,360 CellLink annotations. Abbreviations within annotated text were expanded when Ab3P^48^ identified a corresponding long form in the full text of the article. We identified 14 naming motifs: root, anatomical context, lineage, molecular signature, appearance, functional role, developmental stage, state, variant, molecular signaling, disease, eponym, stimulus, and species/sex. Definitions and examples for each motif are provided in Supplementary Table 8.

All motifs except stimulus were identified through an initial manual review of randomly sampled names. Each name was tokenized and lemmatized using the SciSpacy en_core_sci_sm model^50^. Candidate motif phrases were extracted as contiguous token sequences of length one to three (with overlaps permitted) that did not intersect with a manually labeled motif and did not begin or end with a stop word (e.g., “and”). Candidate phrases were embedded with SapBERT^22^, using the model SapBERT-from-PubMedBERT-fulltext. Candidate motifs were then assigned to names using two complementary approaches:

1. a manually curated mapping from lemma sequences to motif categories, and
2. a multinomial logistic regression classifier trained to predict motif labels from SapBERT embeddings using the manually curated examples as training data.

For each cell population mention, manually labeled motifs were applied first, using the longest non-overlapping matches. All candidate motifs were then passed through the classifier and added in order of increasing margin (difference between the highest and second-highest predicted probabilities).

The manually defined mapping was expanded iteratively, with the classifier used to suggest high-confidence and/or high-frequency candidates for manual review. During this process, we identified the need for a separate stimulus motif to capture references to external sensory cues. Motifs with uncertain meaning or etymology were investigated through searches of PubMed and PMC. In total, approximately 5,000 unique motif phrases were used to label the CellLink annotations; roughly 2,700 were manually labeled by coauthor RL, with the remainder labeled by the classifier. Manually labeled motifs are provided with the source code as described in Code availability. We estimated the accuracy of the classifier to be 0.85±0.02 using a 10-fold stratified cross-validation, repeated five times to reduce variance due to fold assignments.

Lineages were defined by manually mapping high-level CL concepts to a small set of broad categories (e.g., CL:0008019 “mesenchymal cell” → mesenchymal/stromal). Each CellLink annotation was assigned to one or more lineages by propagating its CL identifier to all ancestral CL classes. Mentions whose ancestors spanned multiple lineages were assigned fractionally across lineages. Similarly, related mentions linked to multiple CL identifiers were assigned fractionally across the corresponding lineages. Fractional lineage assignments were propagated to motif counts prior to statistical analysis.

Motif prevalences within a lineage were compared either to the overall prevalence across all lineages or to the corresponding lineage-motif prevalence for the exactly matched cell phenotypes using exact two-sided binomial tests^51^ with any fractional count rounded. Motif prevalences across all lineages were compared using Fisher’s exact test^52^. P-values were corrected for multiple testing with the Benjamini–Hochberg procedure^53^, and differences were considered statistically significant at a false discovery rate (FDR) < 0.01. Counts, p-values and associated q-values are provided in Supplementary File 2.

### Preprocessing of Comparison Corpora

The AnatEM^14^, BioID^15^, CRAFT^16^, and JNLPBA^17^ were downloaded and converted to the BioC XML format^54^. For each corpus, labels corresponding to cell populations or cell types (excluding cell lines) were identified, mapped to the label “cell_type,” and all other annotations were discarded. The label mappings were as follows: AnatEM used “Cell,” BioID used “cell_type,” CRAFT used “CellType,” and JNLPBA used “cell_type.” Identifiers marked as obsolete in CL v2025-01-08 were mapped to their official replacements. We also manually identified and filtered a non-exhaustive set of frequent annotation that conflicted with our guidelines, including overly general terms (e.g., “cells”), named cell lines (e.g., “MCF-7 cells”), cellular components (e.g., “cilia”), and cellular aggregates (e.g. “spheroid”).

## Supporting information

Supplementary figures, tables, note, and algorithm

Annotation guidelines

Cell naming motif data

## Data availability

The CellLink corpus training and validation sets that were developed in the course of this study, have been deposited in Zenodo with the identifier doi:10.5281/zenodo.18090009^55^. To support fair evaluation of cell population NER and EL methods, a redacted version of the CellLink test set is deposited in the same repository. An unredacted version of the CellLink test set is indirectly available to the public via Codabench; users can evaluate their NER and EL methods following the instructions on Codabench at https://www.codabench.org/competitions/13905/.

## Code availability

Computer code that was developed in the course of this study is available in GitHub at https://github.com/noamrotenberg/CellLink-paper.

## Acknowledgements

We thank Alastair Rae for providing the results of the MTIX MeSH indexer.

This research was supported by the Intramural Research Program of the National Institutes of Health (NIH). The contributions of the NIH authors are considered Works of the United States Government. The findings and conclusions presented in this paper are those of the authors and do not necessarily reflect the views of the NIH or the U.S. Department of Health and Human Services. D.O.-S. & C.E.’s research was funded by grant number 2023-221471 from the Chan Zuckerberg Initiative DAF and 220540/Z.20/A, “Wellcome Sanger Institute Quinquennial Review 2021-2026.”

## Author contributions

N.R., R.L., R.I., and R.H.S. designed the research. R.I. designed and guided the annotation process and data validation. H.K., G.T., B.F., and S.R. annotated the article passages. N.R., R.I., R.L., H.K., G.T., B.F., and S.R. wrote the annotation guidelines. R.L. designed and implemented the passage selection algorithm. R.I. and R.L. designed the naming motif analysis, which R.L. implemented. N.R., R.L., and R.I. designed the computational analyses, which N.R. and R.L. performed. D.O.-S. and C.E. performed the chondrocyte case study and edited the Cell Ontology. N.R. and R.L. drafted the manuscript, and all authors contributed to editing. All authors provided discussion and feedback, reviewed the manuscript, and approved the final version.

## Competing interests

The authors declare no competing interests.

## Notes

https://zenodo.org/records/18090009

https://www.codabench.org/competitions/13905/

https://github.com/noamrotenberg/CellLink-paper

### Competing Interest Statement

The authors have declared no competing interest.

