## Supplementary figures, tables, note, and algorithm for "Cell phenotypes in the biomedical literature: a systematic analysis and text mining corpus"

Supplementary materials for “Cell phenotypes in the biomedical literature: a systematic analysis and text mining corpus”

*Affiliations:*
[1]Division of Intramural Research, National Library of Medicine, National Institutes of Health, Bethesda, USA
[2]Division of Immunology, Faculty of Medicine and Health Sciences, and Centre for Bioinformatics and Computational Biology, Faculty of Science, Stellenbosch University, Stellenbosch, South Africa
[3]Wellcome Sanger Institute, Wellcome Genome Campus, Cambridge, UK

[#]These authors contributed equally to this work
[*]Corresponding author

**Table of Contents**

- **Supplementary Table 1**: Medical Subject Headings (MeSH) clusters.
- **Supplementary Table 2**: Comparison of MeSH cluster distributions.
- **Supplementary Table 3**: Illustrative examples of CellLink annotations.
- **Supplementary Table 4**: Inter-annotator agreement by entity type.
- **Supplementary Table 5**: Inter-annotator agreement and number of annotations of each entity type, for each passage type.
- **Supplementary Fig. 1**: Cumulative unique cell population mentions and unique ontology identifiers, as a function of the number of passages annotated over time.
- **Supplementary Table 6**: Descriptive statistics for the CellLink training, validation, and test sets.
- **Supplementary Fig. 2**: Distribution of unique mentions and identifiers across the train-validation-test split.
- **Supplementary Table 7**: Comparison of various statistics on the size and annotations of multiple corpora; expansion of Table 2.
- **Supplementary Fig. 3**: Number of mentions appearing in three frequency ranges across corpora.
- **Supplementary Table 8**: Naming motif definitions and examples.
- **Supplementary Table 9**: Naming motif prevalence for exactly linked cell phenotypes, differentiated by major cell lineage.

- **Supplementary Table 10**: Comparison of naming motif prevalence between mentions labeled as cell phenotype (related) and cell phenotype (exact), differentiated by major cell lineage.
- **Supplementary Table 11**: Comparison of naming motif prevalence between mentions labeled as heterogeneous cell population and cell phenotype (exact), differentiated by major cell lineage.
- **Supplementary Table 12**: Performance of EL models on the CellLink test set (top-k recall).
- **Supplementary Table 13**: Discriminative power of the SapBERT confidence score (AUROC) by entity type.
- **Supplementary Algorithm 1**: Passage Selection Pseudocode.
- **Supplementary Note 1**: Formal definition of evaluation metrics
- **Supplementary Table 14**: Example audit of GPT-5.2 agent performing entity linking.

**Supplementary Table 1 | Medical Subject Headings (MeSH) clusters.** Groups of MeSH terms covering the anatomy of various body systems, diseases of various body systems; also human, mouse, and cell lines. MeSH subheadings are assigned to each MeSH cluster as shown. Not every system had MeSH terms for both anatomical and disease components; these cells are grayed out.

| | **Anatomy Category** | **Disease Category** |
|---|---|---|
| **Blood and Immune** | Hemic and Immune Systems; Erythroid Cells; Myeloid Cells | Hemic and Lymphatic Diseases; Immune System Diseases; Infections |
| **Cancer** | | Neoplasms |
| **Cardiovascular** | Cardiovascular System | Cardiovascular Diseases |
| **Digestive** | Digestive System; Hepatic Stellate Cells; Pancreatic Stellate Cells | Digestive System Diseases |
| **Embryonic** | Embryonic Structures | |
| **Endocrine** | Endocrine System | Endocrine System Diseases |
| **Exocrine** | Exocrine Glands | |
| **Musculoskeletal and Connective** | Musculoskeletal System; Muscle Cells; Chondrocytes; Osteoblasts; Connective Tissue | Musculoskeletal Diseases |
| **Nervous** | Nervous System; Nervous Tissue | Nervous System Diseases |
| **Respiratory** | Respiratory System | Respiratory Tract Diseases |
| **Sensory** | Sense Organs | Eye Diseases; Otorhinolaryngology Diseases |
| **Urogenital** | Urogenital System | Urogenital Diseases |
| **Human** | Humans | |
| **Mouse** | Mice | |
| **Cell lines** | Cells, Cultured; Cell Culture Techniques; Organoids; Culture Media; In Vitro Techniques | |

**Supplementary Table 2 | Comparison of MeSH cluster distributions.** Comparison of articles belonging to the various MeSH clusters in the entire queried subset of PubMed vs. the distribution of the passages in our corpus. Rare MeSH clusters are overrepresented to ensure representation of various biomedical topics. Note that a single article can belong to zero, one, or multiple MeSH clusters, and therefore the columns do not sum to 100%. Not every system had MeSH terms for both anatomical and disease components, these are marked "N/A." Colored shading reflects the range of relative values (min/max), with blue representing higher values and red indicating lower values.

| | PubMed 70k distribution | | CellLink distribution | | Percent of enrichment in CellLink | |
|---|---|---|---|---|---|---|
| | **Anatomy Category** | **Disease Category** | **Anatomy Category** | **Disease Category** | **Anatomy Category** | **Disease Category** |
| **Blood and Immune** | 22.4% | 14.4% | 29.4% | 14.9% | 31.3% | 3.5% |
| **Cancer** | N/A | 25.7% | N/A | 17.8% | N/A | -30.7% |
| **Cardiovascular** | 4.9% | 5.5% | 5.5% | 6.0% | 12.2% | 9.1% |
| **Digestive** | 5.9% | 7.8% | 8.0% | 7.0% | 35.6% | -10.3% |
| **Embryonic** | 4.0% | N/A | 6.5% | N/A | 62.5% | N/A |
| **Endocrine** | 2.3% | 4.1% | 4.2% | 5.7% | 82.6% | 39.0% |
| **Exocrine** | 0.7% | N/A | 3.5% | N/A | 400.0% | N/A |
| **Musculoskeletal and Connective** | 7.5% | 2.9% | 9.0% | 4.9% | 20.0% | 69.0% |
| **Nervous** | 15.3% | 9.8% | 18.3% | 10.0% | 19.6% | 2.0% |
| **Respiratory** | 3.3% | 7.2% | 5.3% | 7.8% | 60.6% | 8.3% |
| **Sensory** | 2.9% | 2.1% | 5.8% | 4.5% | 100.0% | 114.3% |
| **Urogenital** | 4.8% | 6.3% | 6.8% | 7.3% | 41.7% | 15.9% |
| **Human** | 80.0% | | 80.0% | | 0.0% | |
| **Mouse** | 33.9% | | 35.7% | | 5.3% | |
| **Cell lines** | 24.0% | | 19.2% | | -20.0% | |

**Supplementary Table 3 | Illustrative examples of CellLink annotations.**

**A. Frequent, rare (appearing in the corpus only once), and unreported mentions in CellLink, differentiated by entity type.** Abbreviations are explained in brackets outside of quotation marks. Unreported mentions are annotations that did not have an exact match in the Cell Ontology[1] v2025-01-08.

| **Entity type** | **Frequent mentions (frequency in CellLink)** | **Rare mentions** | **Unreported mentions** |
|---|---|---|---|
| Cell phenotype | • 'macrophages', (235)<br>• 'T cells' (197)<br>• 'neurons' (195)<br>• 'endothelial cells' (145)<br>• 'monocytes' (144) | • 'peri-renal or omental adipocytes'<br>• 'scar-associated macrophages'<br>• 'primordial oocytes'<br>• 'Oligodendrocyte precursor cell'<br>• 'ARC neuronal'<br>• 'hepatic NK cells'<br>• 'Meningeal ILC2s' | • 'Activated DCs' [dendritic cells]<br>• 'CD63 + TAMs' [tumor-associated macrophages]<br>• 'atypical memory (AM1 and AM2) B cells'<br>• 'GABAergic Glp1r-expressing neurons' [Glp1r = glycagon-like peptide-1 receptor] |
| Heterogeneous cell population | • 'immune cell' (157)<br>• 'immune cells' (156)<br>• 'PBMCs' (51)<br>• 'tumor cells' (42)<br>• 'PBMC' (37) | • 'leukemic cells'<br>• 'immunosuppressive immune cells'<br>• 'bladder, gallbladder and pancreatic cancer cell'<br>• 'Testis somatic cell'<br>• 'tumor liver vascular endothelial cells'<br>• 'malignant glioma cells'<br>• 'KS spindle cells'<br>• 'hARtg+ prostate tumor cells' | • 'Pancreatic circulating tumor cell'<br>• 'Ocular Surface Epithelial Cells'<br>• 'EpCAM+ CTCs' (circulating tumor cells)<br>• 'PD-1+ immune cells' |
| Vague cell population | • 'immune cell types' (38)<br>• 'immune cell populations' (11)<br>• 'T cell subsets' (11)<br>• 'non-neuronal cells' (8)<br>• 'retinal cell types' (7) | • 'SYMPATHO-ADRENAL CELL TYPES'<br>• 'non-cytotoxic cells'<br>• 'enterochromaffin-like cells'<br>• 'CD326+CD235a- cells'<br>• 'neutrophil sub-clusters'<br>• 'subpopulations of beta cells' | N/A |

**B. Frequent and rare IDs in CellLink, differentiated by entity type.** Vague cell populations are not linked to CL.

| **Entity type** | **Frequent IDs (CL term name, frequency in CellLink)** | **Rare IDs (CL term name)** |
|---|---|---|
| Cell phenotype | • CL:0000235 ('macrophage', 646)<br>• CL:0000084 ('T cell', 622)<br>• CL:0000540 ('neuron', 535)<br>• CL:0000623 ('natural killer cell', 386)<br>• CL:0000576 ('monocyte', 359) | • CL:1001577 ('tonsil squamous cell')<br>• CL:0002491 ('auditory epithelial cell')<br>• CL:0002574 ('stromal cell of pancreas')<br>• CL:0002345 ('CD27-low, CD11b-low immature natural killer cell, mouse')<br>• CL:0002235 ('luminal cell of prostatic acinus')<br>• CL:4052015 ('endocrine gland capillary endothelial cell')<br>• CL:4023011 ('lamp5 GABAergic cortical interneuron')<br>• CL:0008039 ('lower motor neuron')<br>• CL:0000691 ('stellate interneuron') |
| Heterogeneous cell population | • CL:0001064 ('malignant cell', 267)<br>• CL:0001063 ('neoplastic cell', 148)<br>• CL:2000001 ('peripheral blood mononuclear cell', 142)<br>• CL:0000499 ('stromal cell', 71)<br>• CL:0000145 ('professional antigen presenting cell', 36) | • CL:0000167 ('peptide hormone secreting cell')<br>• CL:1000417 ('myoepithelial cell of sweat gland')<br>• CL:0000318 ('sweat secreting cell')<br>• CL:1000448 ('epithelial cell of sweat gland')<br>• CL:1001595 ('rectum glandular cell')<br>• CL:0002142 ('dark cell of eccrine sweat gland')<br>• CL:0000241 ('stratified cuboidal epithelial cell') |

**Supplementary Table 4 | Inter-annotator agreement by entity type.** Vague cell populations are not linked to CL.

| Entity type | Strict span & identifier | Strict span | Approx. span | Strict identifier |
|---|---|---|---|---|
| Cell phenotype | 0.72 | 0.87 | 0.94 | 0.80 |
| Hetero. cell population | 0.52 | 0.58 | 0.60 | 0.57 |
| Vague cell population | N/A | 0.39 | 0.45 | N/A |
| All labels | 0.69 | 0.82 | 0.89 | 0.77 |
| Merged labels* | 0.72 | 0.88 | 0.96 | 0.82 |

* Merged labels refers to combining all annotations into a single type.

**Supplementary Table 5 | Inter-annotator agreement (IAA) and number of annotations of each entity type, for each passage type.** Colored shading reflects the range of IAA values (min/max), with blue representing higher values and red indicating lower values.

| | Inter-annotator agreement* | Cell phenotype | Heterogeneous cell population | Vague cell population |
|---|---|---|---|---|
| **Title** | 0.60 | 141 (75%) | 20 (11%) | 27 (14%) |
| **Section title** | 0.62 | 268 (72%) | 44 (12%) | 59 (16%) |
| **Abstract** | 0.66 | 1,637 (77%) | 187 (9%) | 295 (14%) |
| **Intro** | 0.69 | 3,912 (84%) | 336 (7%) | 384 (8%) |
| **Results** | 0.69 | 5,226 (81%) | 468 (7%) | 738 (11%) |
| **Discussion & Conclusion** | 0.71 | 1,831 (82%) | 157 (7%) | 238 (11%) |
| **Table** | 0.68 | 1,230 (92%) | 73 (5%) | 36 (3%) |
| **Table and figure captions** | 0.69 | 3,988 (84%) | 307 (6%) | 442 (9%) |
| **Other (e.g., abbreviations, appendix)** | 0.70 | 280 (89%) | 24 (8%) | 12 (4%) |

* *Strict span & identifier* agreement across all labels, which requires annotators to exactly match each other's spans, entity type, and identifiers across all entity types. See Methods: Annotation process for further details.

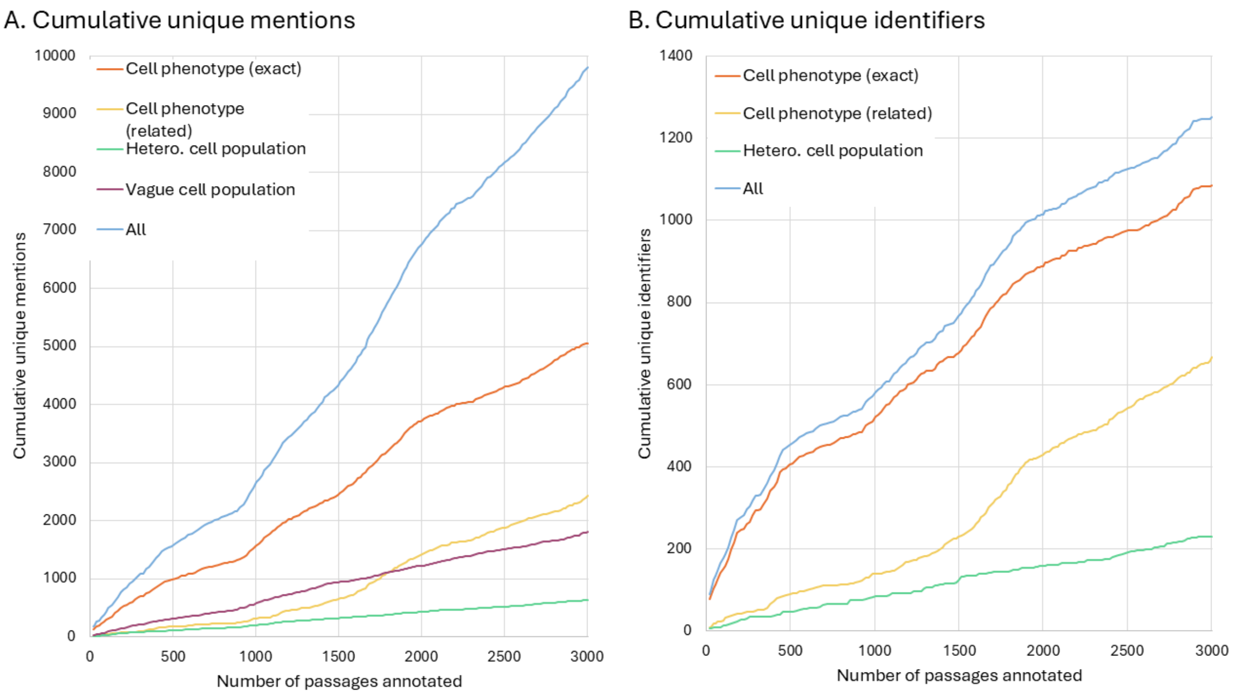


**Supplementary Fig. 1 | Cumulative unique cell population mentions and unique ontology identifiers, as a function of the number of passages annotated over time.** In panel (A), the cumulative number of unique mentions increases approximately linearly across mention types. In panel (B), the cumulative unique identifiers exhibit a more concave trend overall, reflecting a slowing rate of discovery as annotation progresses. Exact matches to CL account for most identifiers and follow the overall trend, while cell phenotypes with related matches to CL and heterogeneous cell populations increase more gradually.  Note that vague cell population mentions were not linked to CL IDs and therefore not included in panel B.

**Supplementary Table 6 | Descriptive statistics for the CellLink training (50.0%), validation (16.7%), and test (33.3%) sets.**

| Statistic | Training | Validation | Test |
|---|---|---|---|
| Tokens | 172,589 | 54,901 | 110,955 |
| Sentences | 7,638 | 2,431 | 4,778 |
| Annotations | 11,140 | 3,591 | 7,631 |
| - Cell phenotype | 9,216 | 3,000 | 6,299 |
| - Hetero cell pop | 811 | 234 | 571 |
| - Vague cell pop | 1,113 | 357 | 761 |
| Unique mentions | 5,410 | 2,203 | 4,007 |
| Unique identifiers | 902 | 603 | 868 |

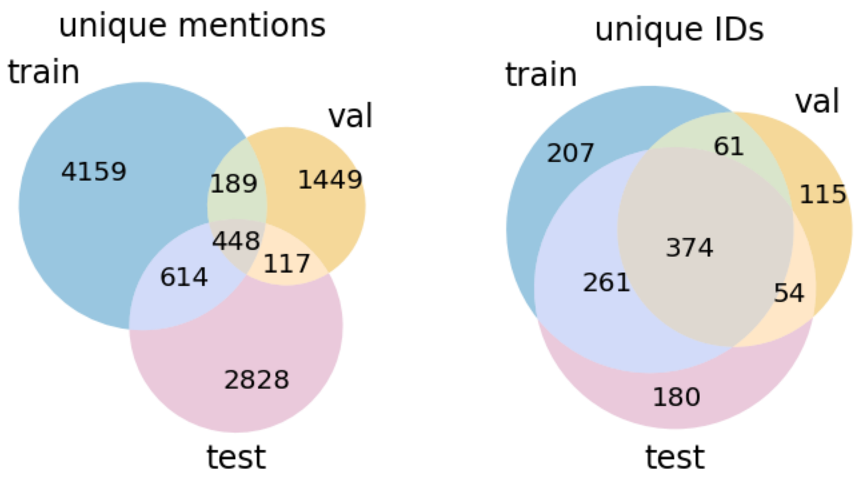


**Supplementary Fig. 2 | Distribution of unique mentions and identifiers across the train-validation-test split.** Venn diagrams showing the overlap in unique mentions and identifiers for the training (train), validation (val), and test sets.

**Supplementary Table 7 | Comparison of various statistics on the size and annotations of multiple corpora (expansion of Table 2).** Comparison corpora were pre-processed to remove non-cell type mentions, see Methods: Preprocessing of Comparison Corpora.

| Statistic | NLM CellLink* | AnatEM[2] | BioID[3] | CRAFT[4] | JNLPBA[5] | SourceData-NLP† | All other corpora |
|---|---|---|---|---|---|---|---|
| Num sentences | 14,847 | 11,285 | 45,449 | 32,284 | 22,437 | 246,766 | 358,221 |
| Num tokens | 338,445 | 259,503 | 771,248 | 652,168 | 564,243 | 4,229,999 | 6,477,161 |
| Num annotations | 22,362 | 3,265 | 4,616 | 5,792 | 8,619 | 22,309 | 44,601 |
| Num annotations per 100 tokens | 6.61 | 1.26 | 0.60 | 0.89 | 1.53 | 0.53 | 0.69 |
| Ann length (mean ± std) | 14.6 ± 10.8 | 13.8 ± 8.4 | 8.3 ± 5.3 | 11.7 ± 7.1 | 16.0 ± 9.7 | 8.0 ± 5.2 | 10.5 ± 7.5 |
| Ann length (min, median, max) | 1, 12.0, 230 | 1, 12.0, 56 | 2, 7.0, 43 | 1, 10.0, 80 | 2, 14.0, 94 | 1, 7.0, 67 | 1, 9.0, 94 |
| Num unique mentions | 9,804 | 1,332 | 610 | 1,028 | 2,765 | 2,052 | 6,502 |
| Unique/total mentions | 0.438 | 0.408 | 0.132 | 0.177 | 0.321 | 0.092 | 0.146 |
| Num unique tokens in annotations | 3,892 | 880 | 470 | 546 | 1,190 | 1,243 | 2,831 |
| Num unique IDs | 1,251 | N/A | 216 | 283 | N/A | 445 | 578 |

* Including *cell phenotype*, *heterogeneous cell population*, and *vague cell population* annotations.

† SourceData-NLP[6] also includes 15,572 annotations labeled "cell_line" and 3,639 unlinked annotations labeled "cell." While all annotations labeled "cell_type" and "cell_line" are linked to identifiers, manual inspection shows that the annotations labeled "cell" include several distinct categories, including specific cell phenotypes (e.g. type II alveolar epithelial cells), cell lines (e.g., CTC-ITB-01) and heterogeneous or vague cell populations (e.g., multinucleated cells, CAR T cells). As including or excluding these annotations would require disambiguation, we excluded SourceData-NLP from further analysis.

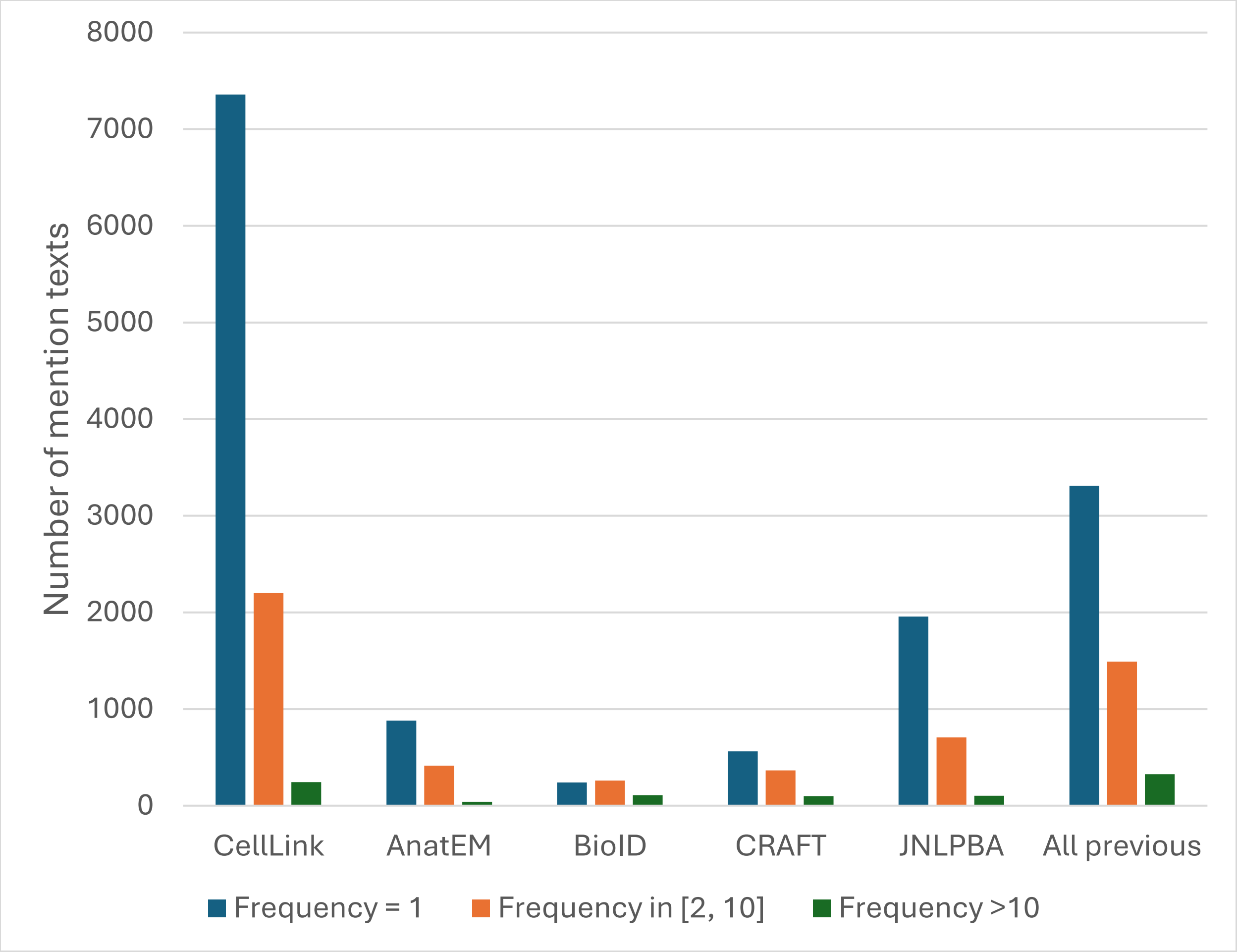


**Supplementary Fig. 3 | Number of mentions appearing in three frequency ranges across corpora.** Comparison of the number of mentions in CellLink and previous corpora, binned into three frequency ranges. “All previous” refers to the union of the previous corpora.

**Supplementary Table 8 | Naming motif definitions and examples.**

| Motif | Definition | Examples |
|---|---|---|
| Root | Fundamental identity nouns corresponding to canonical cell classes, serving as the primary identity term in the name. | neuron, macrophage, fibroblast |
| Anatomical context | Terms that localize the cell within the body, such as tissues, organs, regions or directional descriptors. | vascular, thymic, anterior, retinal |
| Lineage | Terms characterizing developmental origin. | epithelial, mesenchymal, hematopoietic |
| Molecular signature | Terms indicating identifying gene/transcript markers or protein expression patterns. | CD8+, SOX2+, double negative |
| Appearance | Descriptors of visually observable traits such as morphology, structural features, or characteristic staining. | pyramidal, ciliated, acidophilic |
| Functional role | Terms describing biological function or task. | natural killer, suppressor, excitatory, secretory |
| Developmental | Terms indicating the position along a differentiation trajectory, such as maturation stage or lineage commitment. | embryonic, immature, multipotent, terminally differentiated |
| State | Descriptors of dynamic, reversible, or transient physiological conditions. | activated, circulating, exhausted |
| Variant | Labels denoting subtypes or alternative forms within a broader category according to an established classification scheme. | type 1, conventional, non-classical |
| Molecular signaling | Terms describing chemicals used for intercellular communication, including neurotransmitters, hormones, or cytokines. | GABAergic, adrenergic, androgen secreting, calcitonin secreting, histaminergic, interferon-producing |
| Disease | Terms derived from pathological conditions. | tumor-associated, leukemia, rheumatoid, neoplastic |
| Eponym | Names derived from an individual historically associated with the cell type. | Schwann, Purkinje |
| Stimulus | Terms characterizing responsiveness to external physical or chemical stimuli. | photosensitive, NO-sensitive, cold-sensing |
| Species/Sex | Terms indicating origin by organism or biological sex. These terms were not common. | human, mouse, female, male |

**Supplementary Table 9 | Naming motif prevalence for exactly linked cell phenotype mentions, differentiated by cell lineage categories.** Prevalence is defined as the proportion of mentions in which a motif type appears at least once. Values in the “Overall prevalence” column refer to the prevalence for all mentions. Values in all other columns refer to the prevalence difference (prevalence within the specific lineage minus the overall prevalence). Table cells with values not statistically significantly different from the overall prevalence (Benjamini–Hochberg FDR < 0.01) are left blank (see Methods: Naming motif analysis). Colored shading reflects the range of relative values (min/max), with blue representing higher (positive) differences and red indicating lower (negative) differences.

| Motif | Overall prevalence | Epithelial | Endothelial | Mesenchymal/stromal | Muscle | Hematopoietic | Neuronal | Glial | Stem/progenitor | Germ line | Trophoblast/placental | Other |
|---|---|---|---|---|---|---|---|---|---|---|---|---|
| Root | 0.355 | | -0.081 | 0.073 | | | | | | | | |
| Anatomical | 0.142 | 0.101 | 0.143 | 0.098 | 0.166 | -0.083 | | | | -0.123 | | 0.152 |
| Lineage | 0.089 | 0.028 | 0.191 | | -0.069 | -0.035 | -0.061 | | 0.090 | | | |
| Developmental | 0.074 | -0.034 | -0.073 | | | -0.013 | | | 0.181 | 0.144 | | |
| Appearance | 0.059 | 0.025 | | | | -0.014 | 0.062 | | -0.053 | | | |
| Role | 0.055 | -0.039 | -0.053 | -0.042 | | 0.042 | | -0.055 | -0.053 | | | |
| Variant | 0.046 | | | -0.035 | | 0.025 | -0.019 | | -0.042 | | | |
| Molecular signaling | 0.028 | | | | | | 0.060 | | -0.021 | | | |
| Molecular signature | 0.064 | -0.049 | -0.062 | -0.049 | -0.054 | 0.070 | -0.039 | -0.050 | -0.058 | -0.064 | -0.064 | -0.064 |
| Eponym | 0.013 | | | | | -0.006 | | 0.121 | | | | |
| Species | 0.010 | | 0.029 | | | -0.007 | | | 0.015 | 0.052 | | |
| State | 0.036 | -0.028 | -0.031 | -0.023 | | 0.035 | -0.023 | | -0.022 | | | |
| Disease | 0.008 | | | | | | | | | | | |
| Stimulus | 0.001 | | | | | | | | | | | |

**Supplementary Table 10 | Comparison of naming motif prevalence between mentions labeled as cell phenotype (related) and cell phenotype (exact), differentiated by major cell lineage.** Prevalence is defined as the proportion of mentions in which a motif type appears at least once. Values refer to the prevalence difference of the corresponding table cell [cell phenotype (related) minus cell phenotype (exact)]. Table cells with values not statistically significant at FDR < 0.01 are left blank (see Methods: Naming motif analysis). Colored shading reflects the range of values (min/max), with blue indicating higher (positive) differences and red indicating lower (negative) differences.

| Motif | Overall | Epithelial | Endothelial | Mesenchymal/stromal | Muscle | Hematopoietic | Neuronal | Glial | Stem/progenitor | Germ line | Trophoblast/placental | Other |
|---|---|---|---|---|---|---|---|---|---|---|---|---|
| Root | -0.044 | | | | | | -0.060 | | -0.074 | | | |
| Anatomical | | | -0.115 | -0.105 | | 0.017 | | | | | | |
| Lineage | -0.018 | | | | | | | | -0.059 | | | |
| Developmental | | 0.059 | 0.070 | | | | | | | | | |
| Appearance | | | | | | | -0.060 | | 0.023 | | | |
| Role | | | | 0.025 | | -0.022 | | 0.042 | 0.009 | | | |
| Variant | -0.016 | -0.030 | | | | -0.034 | | | | | | |
| Molecular signaling | | | | | | | | | | | | |
| Molecular signature | 0.051 | 0.066 | 0.056 | 0.083 | | 0.039 | 0.060 | 0.043 | 0.056 | | 0.064 | 0.073 |
| Eponym | | | | | | | | -0.093 | | 0.027 | | 0.007 |
| Species | -0.005 | | | | | | | | -0.020 | | | |
| State | 0.015 | | 0.029 | 0.029 | | 0.018 | | | | | | 0.023 |
| Disease | 0.022 | 0.031 | | 0.054 | | 0.030 | | 0.026 | 0.016 | 0.014 | | |
| Stimulus | 0.002 | | | 0.005 | | | 0.010 | | | | | |

**Supplementary Table 11 | Comparison of naming motif prevalence between mentions labeled heterogeneous cell population and cell phenotype (exact), differentiated by major cell lineage.** Prevalence is defined as the proportion of mentions in which a motif type appears at least once. Values refer to the prevalence difference of the corresponding table cell [heterogeneous cell population minus cell phenotype (exact)]. Table cells with values not statistically significant at FDR < 0.01 are left blank (see Methods: Naming motif analysis). Colored shading reflects the range of values, with blue indicating higher (positive) differences and red lower (negative) differences.

| **Motif** | **Overall** | **Epithelial** | **Endothelial** | **Mesenchymal/stromal** | **Muscle** | **Hematopoietic** | **Neuronal** | **Glial** | **Stem/progenitor** | **Germ line** | **Trophoblast/placental** | **Other** |
|---|---|---|---|---|---|---|---|---|---|---|---|---|
| Root | | | | | | | | | | | | -0.082 |
| Anatomical | 0.055 | | | | | 0.118 | 0.178 | | | | | -0.109 |
| Lineage | | | | 0.161 | | | | | | | | |
| Developmental | -0.040 | | | | | | | | | | | |
| Appearance | -0.025 | | | | | 0.063 | | | | | | -0.033 |
| Role | | | | | | | | | | | | -0.050 |
| Variant | -0.037 | | | | | -0.066 | | | | | | |
| Molecular signaling | -0.015 | | | | | | | | | | | |
| Molecular signature | -0.028 | | | | | -0.111 | | | | | | 0.040 |
| Eponym | | | | | | | | | | | | 0.003 |
| Species | | | | | | | | | | | | |
| State | -0.019 | | | | | -0.053 | | | | | | 0.021 |
| Disease | 0.161 | 0.182 | | | | 0.067 | | 0.348 | 0.194 | | | 0.233 |
| Stimulus | | | | | | | | | | | | |

**Supplementary Table 12 | Performance of EL models on the CellLink test set (top-k recall).**
The manually annotated mention spans were used as the input to each model.

| Evaluation | SapBERT | MedCPT-Query | OpenAI text-embedding-3-large |
|---|---|---|---|
| Top-1 | **0.740*** | 0.721 | 0.718 |
| Top-5 | 0.842 | 0.843 | **0.845** |
| Top-10 | 0.882 | 0.888 | **0.900** |

*The highest score for each evaluation is shown in bold font.

**Supplementary Table 13 | Discriminative power of the SapBERT confidence score (AUROC) by entity type.**

| | Cell phenotype | Heterogeneous cell population | Both (labels ignored) |
|---|---|---|---|
| Exact vs. related & no ID | 0.857 | 0.894 | 0.867 |
| Exact & related vs. no ID | 0.831 | 0.704 | 0.850 |

**Supplementary Algorithm 1 | Passage Selection Pseudocode.** Passages were selected by incrementally selecting the passage that would, if added, minimize the Kullback-Liebler[7] (KL) divergence between the set of selected passages and the target distribution. The target distribution is initialized to match the distribution of the input and adjusted according to the configuration to reflect annotation priorities.

```
Inputs: Filtered passages (P); desired number of passages to select (n)

1. Define feature space F as a set of dimensions with corresponding values:
   a. Article-level: journal, year, MeSH terms, MeSH clusters
   b. Passage-level: passage type, passage length, tokens
   c. Semantic-level: PubTator mentions, cell-type mentions from BiomedBERT models
trained on the comparison corpora

2. For each passage p in P:
      Represent p as feature counts across F
      Filter passage if passage type ∈ {'REF/ref','METHODS/paragraph',…}
      Filter passage if length z-score > 4.0
      For dimensions journal, MeSH terms, passage type, tokens:
         Initialize rare feature rate R ← 0.0
         While R < 0.0005: # Expected count is 0.5 per 1000 passages
            Collapse lowest-frequency feature L into R

3. Compute target distribution T over F from P

4. Adjust T:
      a. Adjust for passage length:
         i. Increase: mean length, …
         ii. Decrease: length z-score < -0.2, …
      b. Apply multiplicative scaling:
         i. Increase: MeSH term = 'Single-Cell Analysis', cell-type mentions, …
         ii. Decrease: MeSH cluster = 'Cancer-Disease', cell-line mentions, …
      c. Apply temperature scaling (T = 1.6) to MeSH clusters, tokens, mentions

5. Initialize selected passage set: S ← Ø

6. While |S| < n:
      For each candidate passage c in P:
          For each dimension in F:
             Compute KL divergence between S ∪ {c} and T
      For each dimension in F:
         Z-scale the corresponding KL divergence for each passage
      Sum the z-scaled KL divergence for each passage
      Select c* minimizing divergence
      S ← S ∪ {c*}
      P ← P − {c*}
      If article of c* already has 2 passages:
          Remove all other passages from that article from P

Output: Selected passage set (S)
```

### **Supplementary Note 1 | Formal definition of evaluation metrics.**

*Notation*

Let:

- $G = \{g_1, \ldots, g_m\}$ be the set of reference (gold-standard) annotations.
- $P = \{p_1, \ldots, p_m\}$ be the set of predicted annotations.

Each annotation $a \in G \cup P$ has:

- A document ID identifying the associated passage: $passage\_id(a)$
- A character span $span(a) = [start(a), end(a)]$, where $start(a)$ is the starting character offset (inclusive) and $end(a)$ is the ending character offset (exclusive).
- A type label $type(a) \in \{"cell\ phenotype", "cell\ hetero", "cell\ vague"\}$
- The linked identifier: $identifier(a)$. In general, CellLink annotations may be associated with multiple identifiers, each labeled exact or related. Evaluations for inter-annotator agreement consider any identifier qualified with related to be identical. Evaluations for entity linking ignore both exact and related qualifiers.

Define span overlap as: $overlap(p, g) \Leftrightarrow |span(p) \cap span(g)| > 0$

Precision, recall and F1 score are defined as:

$$Precision = \frac{TP}{TP + FP}, \qquad Recall = \frac{TP}{TP + FN}, \qquad F1 = \frac{2 \cdot Precision \cdot Recall}{Precision + Recall}$$

*Strict NER F1-score*

Define:

- Exact match relation: $M_{strict}(p, g) \Leftrightarrow (passage\_id(p) = passage\_id(g))\ and\ (span(p) = span(g))\ and\ (type(p) = type(g))$
- $TP_{strict} = |(p, g): M_{strict}(p, g)|$
- $FP_{strict} = |P| - TP_{strict}$
- $FN_{strict} = |G| - TP_{strict}$

*Approximate NER F1-score*

Define:

- Eligibility relation: $M_{approx}(p, g) \Leftrightarrow (passage\_id(p) = passage\_id(g)\ and\ overlap(p, g)\ and\ (type(p) = type(g))$
- Let $A \subseteq P \times G$ be a maximum-cardinality matching over all eligible pairs of $M_{approx}$, such that each $p \in P$ and $g \in G$ appears in at most one pair.
  - This matching is computed using the Hungarian algorithm, implemented by `scipy.optimize.linear_sum_assignment`
- $TP_{approx} = |A|$
- $FP_{approx} = |P| - TP_{approx}$
- $FN_{approx} = |G| - TP_{approx}$

*Exact EL F1-score*

Define:

- Exact identifier relation: $M_{EL}(p, g) \Leftrightarrow \ passage\_id(p) = passage\_id(g)\ and\ identifier(p) = identifier(g)$
- $TP_{EL} = |(p, g): M_{EL}(p, g)|$
- $FP_{EL} = |P| - TP_{EL}$
- $FN_{EL} = |G| - TP_{EL}$

*Entity Linking Top-k Recall*

If we let $ID_k(p) = \{identifier_1(p), \dots, identifier_k(p)\}$ be the set of top-k ranked identifiers predicted for a mention, then the corresponding reference annotation is considered correctly predicted if $identifier(g) \in ID_k(p)$.

Top-k recall is defined as:

$$Recall@k = \frac{1}{|G|} \sum_{g \in \mathrm{G}} 1|identifier(g) \in ID_k(p)|$$

**Supplementary Table 14 | Example audit of GPT-5.2[8] agent performing entity linking.**

A. Example agent queries to the Ontology Access Kit[9].

| Mention to link | Agent queries to Ontology Access Kit[*] |
|---|---|
| Myofibroblasts | 1. myofibroblast<br>Agent chose: myofibroblast cell (CL:0000186) |
| Auditory sensory hair cell | 1. auditory sensory hair cell<br>Agent chose: auditory hair cell (CL: 0000202) |
| Short-wavelength cone | 1. short-wavelength cone<br>Agent chose: S cone cell (CL: 0003050) |
| Bipolar cell | 1. bipolar cell<br>Agent chose: ON-bipolar cell (CL:0000749)[†] |
| Condrocyte[‡] | 1. condrocyte<br>2. chondroblast<br>3. chondrocyte<br>4. cartilage cell<br>Agent chose: chondrocyte (CL:0000138) |
| MZB1+ plasma cell | 1. MZB1 plasma cell<br>2. MZB1+ plasma cell<br>3. plasma cell<br>Agent chose: plasma cell (CL:0000786)[§] |
| PV+ cell | 1. PV+ cell<br>2. parvalbumin-positive cell<br>3. Pvalb GABAergic interneuron<br>Agent chose: pvalb GABAergic cortical interneuron (CL:4023018)[‖] |
| SMC[¶] | 1. SMC<br>2. SMC cell<br>3. smooth muscle cell<br>Agent chose: smooth muscle cell (CL:0000192) |

* Note that GPT agent workflows are partially stochastic and may differ given the same input.

† CellLink has this mention with an exact match link to "retinal bipolar neuron" (CL:0000748). When the GPT agent queried the Ontology Access Kit (OAK) for "bipolar cell", "retinal bipolar neuron" was not returned. The GPT agent did not try additional queries to OAK as it was instructed.

‡ The misspelling "condrocyte" was taken directly from the input article.

§ "MZB1+ plasma cell" is not in CL. CellLink has this mention linked as a related match to "plasma cell" (CL:0000786).

‖ While the original article (PMID:32581726) does not explicitly state the meaning of "PV", based on the context, it seems to refer to "parvalbumin" (pvalb). CellLink has this mention linked as an exact match link to "pvalb GABAergic cortical interneuron" (CL:0000786).

¶ "SMC" was annotated as "smooth muscle cell" (CL:0000192) in CellLink. This abbreviation was not expanded by the abbreviation expander.

B. Example agent queries with Ontology Access Kit search results.

| Agent Query to Ontology Access Kit | Response/Ontology Access Kit Search Results |
|---|---|
| `condrocyte` | `"No results were found. Try another query."` |
| `chondroblast` | `[('CL:0000058', 'chondroblast'), ('CL:0007009', 'prechondroblast'), ('CL:0000138', 'chondrocyte')]` |

| | |
|---|---|
| chondrocyte | [('CL:0000138', 'chondrocyte'), ('CL:0000744', 'columnar chondrocyte'), ('CL:0019002', 'tracheobronchial chondrocyte'), ('CL:0020019', 'regulatory chondrocyte'), ('CL:0020020', 'homeostatic chondrocyte'), ('CL:0020022', 'prehypertrophic chondrocyte'), ('CL:0020023', 'effector chondrocyte'), ('CL:1001607', 'articular chondrocyte'), ('CL:0000742', 'periarticular chondrocyte'), ('CL:0000743', 'hypertrophic chondrocyte'), ('CL:1000217', 'growth plate cartilage chondrocyte'), ('CL:2000007', 'articular chondrocyte of knee joint'), ('CL:4072104', 'fibrochondrocyte')] |
| cartilage cell | [('CL:0000138', 'chondrocyte'), ('CL:1000217', 'growth plate cartilage chondrocyte'), ('CL:0000000', 'cell'), ('CL:0007001', 'skeletogenic cell'), ('CL:0002320', 'connective tissue cell'), ('CL:0008019', 'mesenchymal cell'), ('CL:0002639', 'amniotic stem cell'), ('CL:0000459', 'noradrenergic cell'), ('CL:0000499', 'stromal cell'), ('CL:0000586', 'germ cell'), ('CL:0000630', 'supporting cell'), ('CL:0000642', 'folliculostellate cell'), ('CL:0000763', 'myeloid cell'), ('CL:0000988', 'hematopoietic cell'), ('CL:0002242', 'nucleate cell'), ('CL:0002319', 'neural cell'), ('CL:0009004', 'retinal cell'), ('CL:0011026', 'progenitor cell'), ('CL:0011115', 'precursor cell'), ('CL:1001502', 'mitral cell')] |
