## Supplementary material for "Cell phenotypes in the biomedical literature: a systematic analysis and text mining corpus": Annotation guidelines

**Supplementary File 1** for "Cell phenotypes in the biomedical literature: a systematic analysis and text mining corpus"

Noam H. Rotenberg, Robert Leaman, Rezarta Islamaj, Helena Kuivaniemi, Gerard Tromp, Brian Fluharty, Savannah Richardson, Caroline Eastwood, Matthew Diller, Bingfang Xu, Ajith V. Pankajam, David Osumi-Sutherland, Zhiyong Lu, Richard H. Scheuermann

### Annotation Guidelines for the NLM CellLink text corpus

#### Overview

- Our main goal is to identify cell phenotypes. Cell phenotypes include cell types (e.g., fibroblasts, endothelial cells) and their states (e.g., activated, quiescent), if mentioned.
- We also identify two other, less specific, kinds of cell populations, described below: heterogeneous and vague cell populations.
- The annotations produced by this project are intended to be used to create models which can automatically annotate the biomedical literature. We will use these large-scale annotations in downstream analyses, including:
    - Identifying where cell types are discussed in the literature and identifying what cell types are being discussed.
    - Identifying novel cell types not mentioned elsewhere in the literature.
    - Extracting relationships between cell types and other entities, such as diseases.

#### Glossary

- **Cell type:** a classification of cells based on shared features, such as morphology, function, and gene expression
    - We exclude cell lines from this definition!
- **Cell state**: a temporary condition of a cell, e.g., "activated"
- **Cell phenotype**: a distinct, identifiable cell type, including its expressed state, if mentioned
- **Cell population**: a collective term that refers to a group of cells without differentiating them by further; may include cell_type, cell_hetero, and cell_vague
- **Named entity recognition (NER)**: identification of entities in natural language and labeling their span and type
- **Entity linking (EL)**: assigning an identifier to named entities
    - Also known as: named entity normalization (NEN)

- **Relation/Relationship extraction (RE)**: identifying relationships between entities in natural language [out of scope for this project]
- **Span:** an annotated stretch of text, from start to end. The text itself is called the "mention."
    - "approximate span" – refers to a span that overlaps with a ground truth annotation
    - "exact span" – a span that exactly matches the beginning and ending of a ground truth annotation
- **Cell ending**: – common suffixes and "endings" of a cell term
    - e.g., cell, neuron, -cyte, -blast, -clast, -phage
- **Single cell RNA sequencing**: a method of sequencing the RNA of individual cells in a sample; this provides information on the genes expressed by each cell
    - Also known as: scRNA-seq, single cell transcriptomics
    - Similar: single nucleus RNA-seq
    - scRNA-seq experiments commonly identify novel cell types based on distinct patterns of gene expression that can't be differentiated by morphology (e.g., by viewing the population of cells under a microscope)
- **Cluster**: [in the context of scRNA-seq] – a group of cells defined by similarity, often grouped after dimensionality reduction (e.g., UMAP)

### General Guidelines

- Read the assigned text in full (i.e., don't skim).
- Annotation is "open book" (i.e., you may consult the annotation guidelines and internet) but *not* other people, except during collaborative resolution. However, because we are interested in novel cell phenotypes, please keep in mind:
    - Authors may make up their own names; we still want to capture these instances.
    - Authors may discover novel cell types that are not reported anywhere else in the literature.
- Author intent ("library approach"): Annotations represent the statements and apparent meaning of the authors, independent of scientific correctness. Text is annotated as written, even when claims may be incorrect or misleading. Downstream triangulation and confidence assessment will allow extracted information to be filtered later.
- Annotation decisions should be based on the meaning conveyed by the text
    - Annotation decisions should not be made by the surface form alone. Annotators should use the surrounding text from the passage and may use the full article to interpret the intended cell population.
    - Annotation decisions should be made independently of whether a Cell Ontology identifier exists.

- In this document, the example text which is underlined is the span that should be annotated.

#### Annotation Scope

- The primary focus is naturally occurring human and mouse cells.
  - Passages which refer to cell types from other multicellular species (e.g. lamellocytes) should be identified and removed from curation.
- Any bacteria or other unicellular species should not be annotated.
- Cell lines named in the text (e.g., “HeLa,” “Vero cells”) should not be annotated.
  - Populations of cells which are experimentally modified but not named as a cell line should be annotated as described below.
- Anatomical units larger than cells – including tissues – should not be annotated, including organoids
  - E.g., do *not* annotate: “epithelium”, specific types of epithelium, cell cultures, and organoids
- Very broad cell mentions should not be annotated.
  - Do not annotate the word “cell” without further description.
    - E.g., do not annotate “we transfected the cells”
  - Do not annotate highly unspecific mentions: “living cells”, “dead cells”, “cell culture”, “control cells”, “nonnative cells”, “damaged cells”
  - Do not annotate any cell at the organism level or larger.
    - E.g., “human cells”, “mammalian cells”, “patient-derived cells”
  - Do not annotate experiment-specific descriptors, which do not provide any information about a cell population outside of the experiment.
    - E.g., “td-Tomato+ cells”, “GFP+ glial cells” (only annotate “glial cells”), “cluster 4 hepatocytes” (only annotate “hepatocytes”)

#### Cell Population Type Definitions

We have three categories of cell populations:

- ***Cell phenotype***; label: `cell_phenotype`
  - Definition: An unambiguously distinct, identifiable cell type, including its expressed state, if mentioned
  - These groups of cells would be expected to share a transcriptomic profile – a pattern of gene expression that allows them to be distinguished from other cell populations.
  - E.g., “hepatocytes,” “microglia”, “activated fibroblast”

- ***Heterogeneous cell population***; label: `cell_hetero`
  - Definition: A group of cells from unrelated lineages, typically characterized by a shared attribute
    - These cells are grouped by an attribute that does not define the phenotype.
    - e.g., "kidney cells", "secreting cells", "progenitor cells", "BCC (Basal Cell Carcinoma) cells"
    - These are sometimes very broad, e.g., "BCC cells"
  - Almost always, a group of cells described only by their source tissue or organ is cell_hetero. E.g., "kidney cells", "retinal pigment epithelium cells"
    - Example exception: authors sometimes give a full description initially but drop some details later that are clear from context; "lung cells" might therefore refer to "lung epithelial cells" if the "epithelial" aspect is clear in context and would be annotated as cell_phenotype.
  - Note: these annotations are heterogeneous, and they may be broad, but they are *not* vague, e.g., "microglia-like cells" is a vague cell population (i.e., cell_vague, *not* cell_hetero)
  - Heuristic: A population of cells that is described solely in terms of anatomical context should likely be classified as cell_hetero (e.g., "renal cell") but adding an anatomical description to a cell phenotype would likely still be classified as cell_phenotype (e.g., "T cell" is a cell_phenotype, so "lung T cell" is also a cell_phenotype).
- ***Vague cell population***; label: `cell_vague`
  - Definition: A group of cells whose identities cannot be determined from the text provided
  - They are often described (rather than named) because there is no name.
    - Consequently, this group has a highly variable linguistic structure.
  - A common feature of this group is that additional context would be needed to reproduce the cells type.
    - e.g., "Ly6a, Cebpb, and Egfr expressing cells" [the markers don't define the cell], "microglia-like cells"
      - Exception: "hillock-like cells" is a cell_phenotype because it has become an established name.
      - Counter-example: "CD8+ cells" is a cell_phenotype in most contexts, because it clearly refers to CD8+ T cells in most contexts.
    - This group includes experimental conditions and experiment-specific cells.
      - E.g., "HTNV-exposed primary monocytes", "CAR T cells"
  - Heuristic: Cell types that start with "non" are generally descriptions and should be annotated as vague cell populations, e.g., "<u>non-kidney cell</u> contaminants"

#### Span Selection Guidelines

- Basic principle: annotate the shortest span necessary to encapsulate the entire, distinct cell population, plus the cell ending (if it is present).
  - Encapsulating distinct cell phenotype: e.g., "activated CD8+ MAIT cells" – each word in this span is essential to distinguishing the specific cell phenotype
  - Cell ending examples:
    - Annotate "fibroblast cells" even though technically "fibroblast" could be a cell type on its own
    - Annotate "cone photoreceptor cell" even though technically "cone", "photoreceptor", "cone photoreceptor", and "photoreceptor cell" could each be a cell type on their own
    - Examples when the cell ending is not available: see "cell ellipsis"
- For a given span, only annotate the single most specific cell population being discussed. Do not create additional annotations for nested or partial cell population mentions within the same span.
  - E.g., annotate "activated CD8+ MAIT cells", but do not create additional annotations for "CD8+ MAIT cells" and "MAIT cells"
  - Exception: a cell population described with a **subset descriptor** should follow the rules for Subset descriptors described below.
- Annotate entire words, not portions of words, unless necessary.
  - E.g., annotate "peripheral blood mononuclear cells (PBMCs)" [do not leave the "s" out of the annotations]
  - Counter example: the paper has a typo, and there is no space between a cell type and an unrelated word. Annotate the cell type as the author intended it.
- Modifiers to include in the span:
  - Include marker genes, even if redundant.
    - Particularly in the case of CD68+ macrophages; even though CD68 is a marker for macrophages, a paper can and will discuss CD68– macrophages
  - If a cell subtype is defined by gene expression or any other descriptive word, then it should be included in the span, e.g., "CD8+ T cells" is distinct from T cells; "mucosal-invariant T cells" is distinct from T cells
    - Always include genes used as descriptors.
  - By default, any descriptor implies a distinct subtype, unless it is clearly redundant (without needing specialized knowledge in a specific subset of cell biology)
    - E.g., "tumor-infiltrating dendritic cells"

        - E.g., do *not* stretch a span to include phrases which are clearly redundant such as "healthy," "adult," "mature," and "terminally differentiated" [see info on shortest complete span]
            - For example, a "mature monocyte" is exactly a monocyte
            - In some cases, "mature" and "terminally differentiated" should be annotated because the phrase distinguishes the annotation from the base cell phenotype, e.g., "terminally differentiated B lineage cell" is a B cell, which is a specific subtype of B lineage cell
            - Exception: annotate the entire span of "normal and tumor mesothelial cells" because they are actually a coordination ellipsis of 2 different cell types
        - "CD8-expressing T cell" is a cell phenotype; we consider it to be a name, equivalent to "CD8+ T cell"
- Modifiers to exclude:
    - Don't include "human" (or other species), unless it is required, e.g., the species name is part of an abbreviation.
        - E.g., annotate: "human umbilical vascular endothelial cells (HUVEC)", "mouse embryonic fibroblasts (MEF)"
    - Don't annotate "primary" as it relates to cell passaging (unless required for another reason in the guidelines).
        - Example of an exception: "HTNV-exposed primary monocytes"; "primary" is annotated in order to annotate other essential descriptors.
- Coordination ellipsis – when "and" (or another conjunction) carries over a portion of a cell population name
    - In these cases, annotate the entire span necessary to completely encapsulate all of these cell populations into 1 annotation
        - E.g., "we observed the immune and endothelial cells" implies "we observed the immune cells and endothelial cells"
        - E.g., "we observed the rod and cone cells" – in this case, "rod" could technically stand on its own but that is not the author's intention (i.e., the author did not intend to state "we observed the cone cells and the rod")
    - Our annotation tool only allows one label per span, so even though coordination ellipses refer to more than one entity, it can only be labeled with one entity type.
        - A coordination ellipsis with cell_phenotype & cell_hetero instances is cell_hetero.
        - A coordination ellipsis that includes a cell_vague is always cell_vague.
- Subset descriptors – descriptors that refer to an unspecified subset of the base cell type.
    - Unspecified subtypes are *not* cell phenotypes or cell_hetero, but they are definitely referring to a population of cells; therefore, they should be marked as cell_vague

- Common subset descriptors: “clusters”, “subtypes”, “population”, “subpopulation”, “classes”, “types”, “subset”, “phenotype”
- E.g.,:
  - Annotate “glial cluster” as “glial” (cell_phenotype [subset ellipsis]) and “glial cluster” (cell_vague)
  - Annotate “ciliated cell types” as “ciliated cell” (cell_hetero) and “ciliated cell types” (cell_vague)

### Additional Rules

- Acronyms: annotate acronyms and abbreviations as if they were completely written out
  - E.g., “The human <u>peripheral blood mononuclear cells</u> (<u>PBMCs</u>) were…”
  - If an acronym interrupts another span, then include it in that span
    - E.g., annotate “<u>mucosal-associated invariant T (MAIT) cells</u>” as a single cell_phenotype annotation
- Cell population mentions within other nouns:
  - Annotate cell population mentions, even when it is a subpart of a larger noun chunk (i.e., it is mentioned within another noun).
    - We do this because the larger noun is referencing the cell population.
  - E.g., “Chinese hamster <u>ovarian cell</u> line (CHOC)” [annotation for “ovarian cell” is cell_hetero]; do not annotate “CHOC” because it is a cell line
- Adjective forms
  - The adjective form of a cell population (e.g., "phagocytic" from phagocyte) is annotated based on the meaning of the overall phrase
  - If the phrase definitely refers to a cell population, but the word “cell” or other cell ending is dropped, then the adjective is annotated (this is a “cell ellipsis”).
    - e.g., “glial clusters”, “endothelial markers”. In these cases, annotate “glial” and “endothelial” as cell_phenotype because they are definitely referring to glia / glial cells and endothelial cells, respectively.
    - Analyze the authors’ intentions.
    - Heuristic: If you can add the word “cell” without changing the meaning of the sentence, then that is a cell ellipsis.
  - If the phrase uses the adjective form to refer to some aspect of the cell besides a cell population, then the adjective is not annotated
    - E.g., annotate “myeloid” in “<u>myeloid</u> differentiation” but not “myeloid oncogene”
- Diseased cells
  - Cell populations that share a phenotype should be annotated as a cell phenotype, even if the phenotype only occurs in the context of disease. E.g., “cancer-associated fibroblast”

    - Tumor or cancerous cells are cell_hetero. While all cancer cells derive from a single cell, we consider that they have become far enough apart. E.g., “Basal cell carcinoma cells”
- Span overlapping:
    - Novel and/or vague cell populations are frequently described using existing cell types
        - A cell_vague span can contain both cell_phenotype and cell_hetero
        - E.g., annotate “microglia-like cells” as two annotations: “microglia” (cell_phenotype) and “microglia-like cells” (cell_vague)
        - In these cases, the cell_vague span should completely contain the other annotation, not partially overlap.
    - Other overlaps are not allowed
        - cell_phenotype and cell_hetero annotations cannot overlap because cell_phenotype and cell_hetero are both specific populations, and their spans won’t be intermingled unless there is an ellipsis (which is a single annotation; see guidelines on ellipses).
        - An entity cannot have overlapping annotations of the same entity type; cell_phenotype annotations cannot overlap with another cell_phenotype annotation
- Specific cases:
    - “immune cells” are cell_hetero because there are multiple lineages (yolk sac and hematopoietic)
    - Named T cell populations (e.g., memory T cell, effector T cell) should be annotated as cell_phenotype unless specifically stated to be from heterogeneous lineages.

### Identifier Selection Guidelines

- Eligibility
    - Most cell_phenotype and many cell_hetero annotations can be linked to an identifier in Cell Ontology (https://www.ebi.ac.uk/ols4/ontologies/cl)
    - cell_vague cannot be linked – their identity is unclear, by definition.
- Context & author intent
    - When assigning identifiers to a span, use the context to assign an identifier that more accurately captures the intention of the author.
        - E.g., if an author is referring to cardiac fibroblast cells using the phrase “fibroblast cells,” then link “fibroblast cells” to CL:0002548 (“fibroblast of cardiac tissue”)

    - If the author implies there is a distinction, then we trust the author (because we have no other way to know; we are extracting the information reported in the literature)
    - E.g., we assume that "pectoral muscle motor neurons" are distinct from other motor neurons unless we have evidence otherwise; same with "proliferating endothelial cells" vs. endothelial cells
- Matching degrees
    - Search for cell_phenotype and cell_hetero in the Cell Ontology using the Ontology Lookup Service: [https://www.ebi.ac.uk/ols4/ontologies/cl](https://www.ebi.ac.uk/ols4/ontologies/cl)
    - Any phrase listed as "synonyms" is considered equivalent to the name of the cell phenotype.
    - If there is no Cell Ontology term or synonym with the same meaning:
        - If a similar name is available, then use judgement whether the annotated span is distinct from the similar name
        - Anything reproducibly distinct should not be considered an exact match (e.g., activated MAIT cell is distinct from a MAIT cell)
    - If the meaning of the phrase and term are the same; mark it as an exact match by starting the identifier with the prefix "(skos:exact)"
    - If no exact match is available, then look for a related match; mark related identifiers with the prefix "(skos:related)"
        - Related matches can be the nearest parent, or sister/cousin cell type, whichever is the best fit.
        - If there are multiple related matches that you believe are very relevant, you can add multiple identifiers, comma-separated.
        - For cell_hetero, you can add both the nearest parent/sister/cousin cell type in addition to enumerating any specific cell phenotype that is included in the heterogeneous group of cells.
        - If you cannot find a related match, leave the identifier blank (or as a dash in the case of coordination ellipsis)
- Specific scenarios
    - When a single annotation includes more than one cell phenotype (because of a coordination ellipsis), use semicolons to separate identifiers (no space).
        - Write the identifiers in the order that their corresponding mentions appear.
        - If one or more of the cell types don't have an identifier, then put a dash in its place
    - By default, any gene or descriptor implies a distinct subtype, unless it is clearly redundant (without needing specialized knowledge in a specific subset of cell biology)
    - Diseased cells and disease-associated cells

- There are few if any diseased cells in the Cell Ontology; therefore, any diseased cells cannot be linked to a CL term with an exact match, though they may be linked as related (skos:related).
- Cells which are cancerous may be linked (skos:related) to both the cell type of origin and term for malignant cells.
    - E.g.: “glioblastoma cells” may be linked to “glial cells” (CL:000030) and “malignant cell” (CL:0001064)
- Healthy cells that are associated with cancer or tumors should not be linked to the cancer or tumor CL ID, because these cells are not cancerous or neoplastic themselves. E.g., “cancer-associated fibroblast”, “tumor-infiltrating lymphocyte”
